# Glucosylhydroxyceramides Modulate Secretion Machinery of a Subset of Plasmodesmata Proteins and a Change in the Callose Accumulation

**DOI:** 10.1101/2020.03.31.017558

**Authors:** Arya Bagus Boedi Iswanto, Jong Cheol Shon, Minh Huy Vu, Ritesh Kumar, Kwang Hyeon Liu, Jae-Yean Kim

## Abstract

The plasma membranes encapsulated in the plasmodesmata (PDs) with symplasmic nano-channels contain abundant lipid rafts, which are enriched by sphingolipids and sterols. The attenuation of sterol compositions has demonstrated the role played by lipid raft integrity in the intercellular trafficking of glycosylphosphatidylinositol (GPI)-anchored PD proteins, particularly affecting in the callose enhancement. The presence of callose at PD is tightly attributed to the callose metabolic enzymes, callose synthases (CalSs) and β-1,3-glucanases (BGs) in regulating callose accumulation and callose degradation, respectively. Sphingolipids have been implicated in signaling and membrane protein trafficking, however the underlying processes linking sphingolipid compositions to the control of symplasmic apertures remain unknown. A wide variety of sphingolipids in plants prompts us to investigate which sphingolipid molecules are important in regulating symplasmic apertures. Here, we demonstrate that perturbations of sphingolipid metabolism by introducing several potential sphingolipid (SL) pathway inhibitors and genetically modifying SL contents from two independent SL pathway mutants are able to modulate callose deposition to control symplasmic connectivity. Our data from pharmacological and genetic approaches show that the alteration in glucosylhydroxyceramides (GlcHCers) particularly disturb the secretory machinery for GPI-anchored PdBG2 protein, resulting in an over accumulated callose. Moreover, our results reveal that SL-enriched lipid rafts link symplasmic channeling to PD callose homeostasis by controlling the targeting of GPI-anchored PdBG2. This study elevates our understanding of the molecular linkage underlying intracellular trafficking and precise targeting to specific destination of GPI-anchored PD proteins incorporated with GlcHCers contents.

## Introduction

Plasmodesmata (PDs), plant-specific symplasmic channels, cross the plant cell wall and physically connect the cytoplasm and endoplasmic reticulum (ER) of contiguous cells. These intercellular channels play important roles in multicellular events during plant development by allowing the molecular exchange of signaling molecules such as transcription factors, RNAs, and growth regulators (Zambryski and Crawford, 2000; Maule, 2008; Wu et al., 2018). The plasmodesmal plasma membrane (PD-PM) is distinct from general cellular plasma membrane (PM) and is characterized by the enrichment of sterols and sphingolipid SL species (Grison et al., 2015; Iswanto and Kim, 2017; Mamode Cassim et al., 2019). The domains enriched with those special lipids are referred to as the membrane microdomain compartments, also called lipid rafts (Mongrand et al., 2010).

The dynamic regulation of size exclusion limit (SEL) of PDs particularly involves the modulation of callose deposition; callose is synthesized by callose synthases (CalSs) and degraded by β-1,3-glucanases (BGs) (Verma and Hong, 2001; Levy et al., 2007; Chen and Kim, 2009). The importance of sterols in the function of PDs has well been reported; the perturbation of sterol biosynthesis pathways affects symplasmic intercellular connectivity by changing the subcellular localization of glycosylphosphatidylinositol (GPI)-anchored PD proteins, namely, plasmodesmata callose binding-1 (PDCB1) and plasmodesmata-associated β-1,3-glucanase (PdBG2), which is followed by callose accumulation (Grison et al., 2015). Recently, by characterizing PHLOEM UNLOADING MODULATOR encoding a putative enzyme required for biosynthesis of SLs with very-long-chain fatty acid, SLs were shown to modulate plasmodesmal ultrastructure and phloem unloading (Yan et al., 2019). In addition, an increase in ceramide content resulted in a reduced the callose deposition in Arabidopsis cell (Bi et al., 2014). However, the involvement of sphingolipids (SLs) in callose-mediated PD regulation has not yet been established.

GPI lipid anchoring refers to the post-translational modification of many cell-surface proteins with a glycolipid. Lipid rafts provide platforms for GPI-anchored proteins, enabling them to stay at the cellular membrane (Mongrand et al., 2010). Modulations in lipid raft components have been implicated in the secretory machinery affecting certain PM proteins, in which the ABCB19 and PIN2 translocations are perturbed along with the depletion of sterols and SLs (Yang et al., 2013; Wattelet-Boyer et al., 2016). GPI-anchored PD proteins contain a signal peptide (SP) at their N-terminal domain, and their GPI modification occurs in the ER. Attachment of the GPI moiety to the carboxyl terminus (omega-site) of the polypeptide occurs after proteolytic cleavage of a C-terminal propeptide from the proprotein (Udenfriend and Kodukula, 1995). Moreover, the GPI anchor undergoes remodeling either in the glucan chain or in the lipid moiety by the addition of a saturated fatty acid chain. GPI remodeling is crucial for the attachment events of GPI-anchored proteins to the membrane lipid raft during translocation (Muniz and Zurzolo, 2014; Kinoshita, 2015).

The SL synthetic pathway is initiated by the activation of serine palmitoyl transferase, which is involved in the conversion of serine and palmitoyl-coA to 3-ketosphinganine; 3-ketosphinganine then proceeds to form ceramides as intermediate products in the metabolism of SL (Michaelson et al., 2016). Ceramide undergoes modification to produce glycosphingolipids, and glycosylinositolphosphorylceramides (GIPCs) and glucosylceramides (GlcCers) are two major classes of glycosphingolipids found in plants (Warnecke and Heinz, 2003; Kim et al., 2013). Approximately 64% of all SLs are GIPCs, and several studies have shown that critical roles played by GlcCers and GIPCs in intracellular trafficking and plant developmental events (Markham and Jaworski, 2007; Msanne et al., 2015; Fang et al., 2016). Furthermore, GlcCers can be divided into 6 groups with distinct molecular compositions depending upon the existence and attributes of types of long-chain-bases (LCBs), double bond and hydroxy fatty acids (Hamanaka et al., 2002).

Here, using pharmacological and genetic approaches we show that the perturbation of SL metabolic pathways results in the alteration of symplasmic connectivity in a manner that is linked to the PD callose level. In addition, we found that the attenuation of GlcHCers alters the subcellular localization of GPI-anchored PdBG2 and PDCB1 proteins, not for non GPI-anchored PDLP1 protein. We also report that there are at least two independent secretory pathways for GPI-anchored PD protein and non-GPI-anchored PD protein. In summary, our studies reveal that GlcHCers are important for the regulation of symplasmic channeling through the modulation of PD targeting of GPI-anchored PdBG2 protein and the callose level.

## RESULTS

### Perturbation in SL metabolism alters PD permeability in Arabidopsis hypocotyl and root tip

To test if SLs are important in PD regulation, we used several potential inhibitors of SL metabolism (Edsall et al., 1998; Singh et al., 2000; Coursol et al., 2003; Sperling and Heinz, 2003; Wright et al., 2003; Falcone et al., 2004; Chen et al., 2006; Delgado et al., 2006; Kang et al., 2008; Worrall et al., 2008; Melser et al., 2010; Markham et al., 2011; Su et al., 2011; Adibhatla et al., 2012; Chen et al., 2014; Michaelson et al., 2016) (**Supplemental Fig. S1**). We performed PD permeability assays using 8-hydroxypyrene-1,3,6-trisulfonic acid (HPTS) (Han et al., 2014) and aniline blue-callose staining assays with etiolated Arabidopsis hypocotyls after treatment with SL synthetic pathway inhibitors. PD permeability, as shown by HPTS movement, was reduced in seedlings treated with myriocin, FB1, PDMP, and D609, but seedlings treated with DMS displayed enhanced PD permeability (**Supplemental Fig. S2**). Accordingly, PD permeability was inversely associated with callose deposition; higher PD permeability was found at a lower level of callose deposition (**Fig. 1 A1-A6, B**). Furthermore, we also investigated the action of SL synthetic pathway inhibitors in PD regulation at Arabidopsis root tip. At 24 h after the treatment of SL inhibitors such as myriocin, FB1, PDMP and D609 the callose levels were significantly increased as compared with mock condition, whereas DMS treatment reduced the callose level (**Fig. 1 C1-C6, D**).

**Figure 1.**
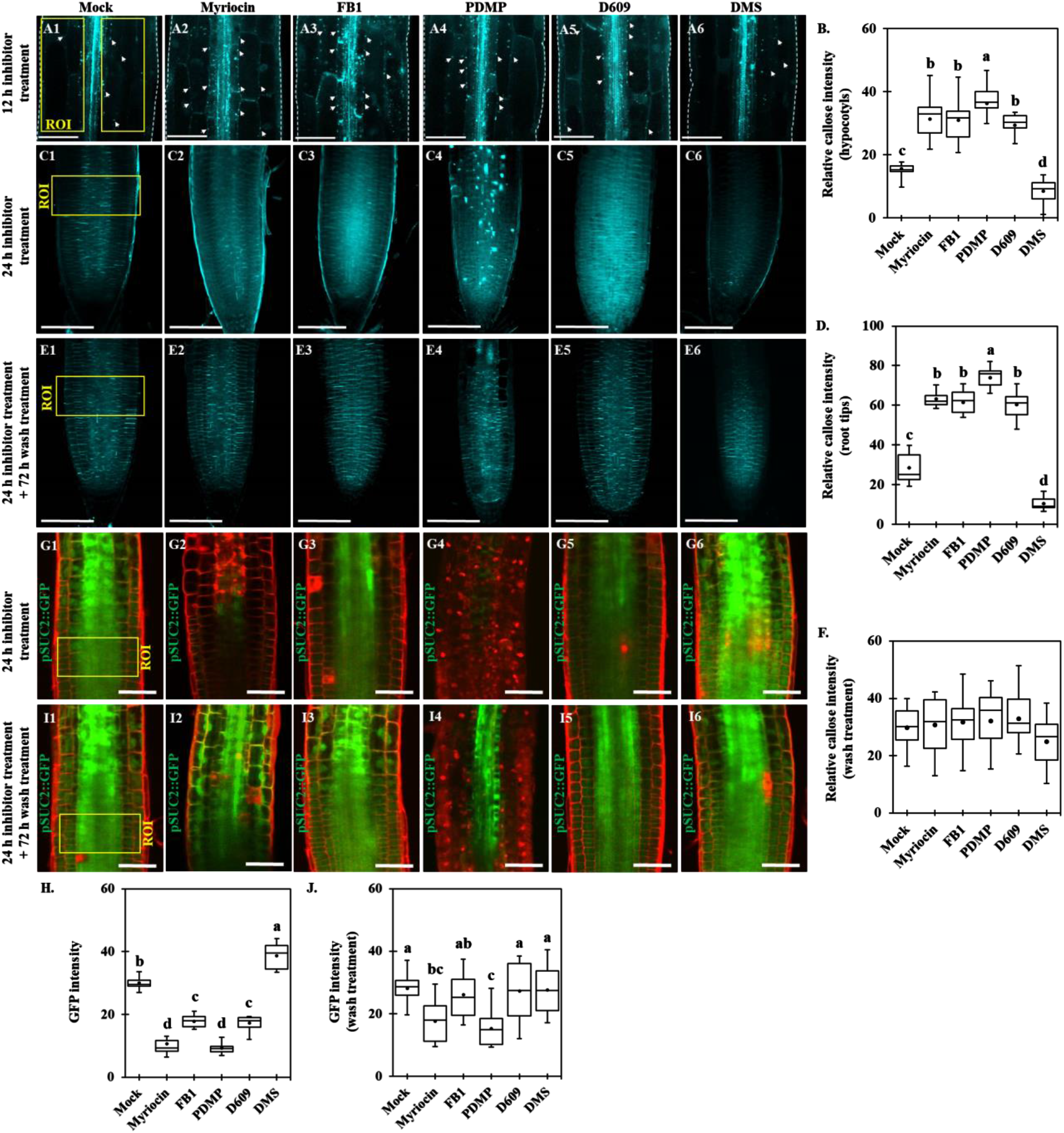
Perturbation in the SL pathway alters PD permeability in Arabidopsis hypocotyls and root tips. (**A1-A6**) Confocal images show aniline blue-stained callose in Arabidopsis etiolated hypocotyls after SL inhibitor treatment (12 h). White arrows show aniline blue-stained callose at PD. Scale bars = 100 μm. (**B**) Quantitative data show the relative callose intensity of Arabidopsis etiolated hypocotyls after SL inhibitor treatment (**A1-A6**) (n = 15). (**C1-C6**) Confocal images show aniline blue-stained callose in Arabidopsis root tips after SL inhibitor treatment (24 h). White arrows show aniline blue-stained callose at PD. Scale bars = 100 μm. (**D**) Quantitative data show the relative callose intensity of Arabidopsis root tips after SL inhibitor treatment (**C1-C6**) (n = 15). (**E1-E6**) Confocal images show aniline blue-stained callose in Arabidopsis root tips. Images were captured right after wash treatment (replaced to the normal agar plates, 72 h) from SL inhibitor treatments (24 h). Scale bars = 100 μm. (**F**) Quantitative data show the relative callose intensity of Arabidopsis root tips after wash treatment (**E1-E6**) (n = 9). (**G1-G6**) Confocal images of root tips of Arabidopsis transgenic plants expressing pSUC2::GFP. Images were captured right after SL inhibitor treatment (24 h). Scale bars = 50 μm. (**H**) Quantitative data show the GFP fluorescence intensity of Arabidopsis root tips expressing pSUC2::GFP after SL inhibitor treatments (**G1-G6**) (n = 12). (**I1-I6**) Confocal images of root tips of Arabidopsis transgenic plants expressing pSUC2::GFP. Images were captured right after wash treatment from SL inhibitor treatment (24 h). Scale bars = 50 μm. (**J**) Quantitative data show the GFP fluorescence intensity of Arabidopsis root tips expressing pSUC2::GFP after wash treatment (**I1-I6**) (n = 12). Yellow boxes indicate region of interest (ROI) for callose quantification (**A1-A6, C1-C6**) and GFP fluorescence intensity quantification (G1-G6, I1-I6). All 3 independent biological experiments were performed and statistical significances were done by One-Way ANOVA with Tuckey-Kramer test. The black dots and lines in boxes (**B, D, F, H, J**) indicate the mean and median, respectively.

Next, we also used a transgenic plant expressing GFP under the control of a companion cell-specific AtSUC2 promoter (pSUC2::GFP) (Benitez-Alfonso et al., 2013) to test if the symplasmic diffusion of GFP can be affected by SL pathway inhibitors-altered callose accumulation. The GFP fluorescence intensity at the root tip was observed 24 h after mock or inhibitor treatment. The GFP fluorescence intensities in seedlings treated with myriocin, FB1, PDMP and D609 were significantly reduced compared with the GFP fluorescence intensity in seedlings subjected to the mock treatment. In contrast, DMS treatment resulted in enhanced PD permeability, as shown by the higher intensity of the GFP fluorescence (**Fig. 1 G1-G6, H**). Although we cannot exclude a possibility that some SL pathway inhibitors also altered pSUC2-mediated gene expression or protein synthesis, in general, GFP fluorescence intensities in the root meristems were inversely associated with callose deposition. We next tested whether the effect of SL inhibitor treatment on callose level and GFP movement were reversible. When SL inhibitor-treated seedlings were placed back to Murashige and Skoog (MS) control medium for 72 h, we observed no obvious differences between callose levels (**Fig. 1 E1-E6, F**) as well as GFP fluorescence intensities (**Fig. 1 I1-I6, J**) in mock condition and SL inhibitor-treated seedlings. These results strongly suggest that perturbation of SL metabolic pathways alters PD permeability by modulating callose accumulation.

### GlcHCers are important for GPI-anchored PD protein localization

To explore the role of SL metabolites in the regulation of PD permeability, we performed SL profile analyses in 14-day-old Arabidopsis seedlings after a 24 h inhibitor treatment. After myriocin treatment, the total contents of SLs and individual contents of several SL molecules such as LCBs, GlcCers, GlcHCers and GIPCs were strongly reduced (**Fig. 2A, D-F, I-K**). FB1-treated seedlings displayed reductions in the total contents of SLs and several SL molecules such as GlcCers, GlcHCers, and GIPCs (**Fig. 2D-F, I-K**), but the total contents of LCBs and ceramides were significantly increased (**Fig. 2A, B**). However, several ceramide molecules with very long-chain fatty acids (VLCFAs) were significantly reduced, conversely ceramide molecules with long-chain fatty acids (FCLAs) such as d18:0/16:0, t18:1/16:0 and t18:1/16:0 were highly elevated (**Fig. 2G**). PDMP-treated seedlings resulted in the attenuation of total contents of GlcCers, GlcHCers and GIPCs (**Fig. 2D-F**), but the total contents of SLs and several SL molecules such as LCBs, ceramides and hydroxyceramides were elevated (**Fig. 2A-C, G, H**). Moreover, after D609 treatment, the total contents of GIPCs and GlcHCers were significantly reduced (**Fig. 2E, F**), but the total contents and several SL molecules of LCBs, ceramides and hydroxyceramides were significantly elevated (**Fig. 2A-C, G, H**). Next, DMS-treated seedlings resulted in the enhancement of the total contents of hydroxyceramides, GlcHCers and GIPCs without disturbing the other SL molecules levels (**Fig. 2C, E, F**). Based on the callose phenotype, 4 SL inhibitor treatment (myriocin, FB1, PDMP and D609) showed callose accumulation enhancement, whereas DMS treatment showed the reversed result. When we observed the SL results, we found that myriocin-, FB1-, PDMP- or D609-treated seedlings similarly resulted in the attenuation of GlcHCers level, conversely, the level of GlcHCers was elevated in the DMS treatment. Altogether, our results suggested that GlcHCers level is important for the alteration of callose-mediated PD permeability.

**Figure 2.**
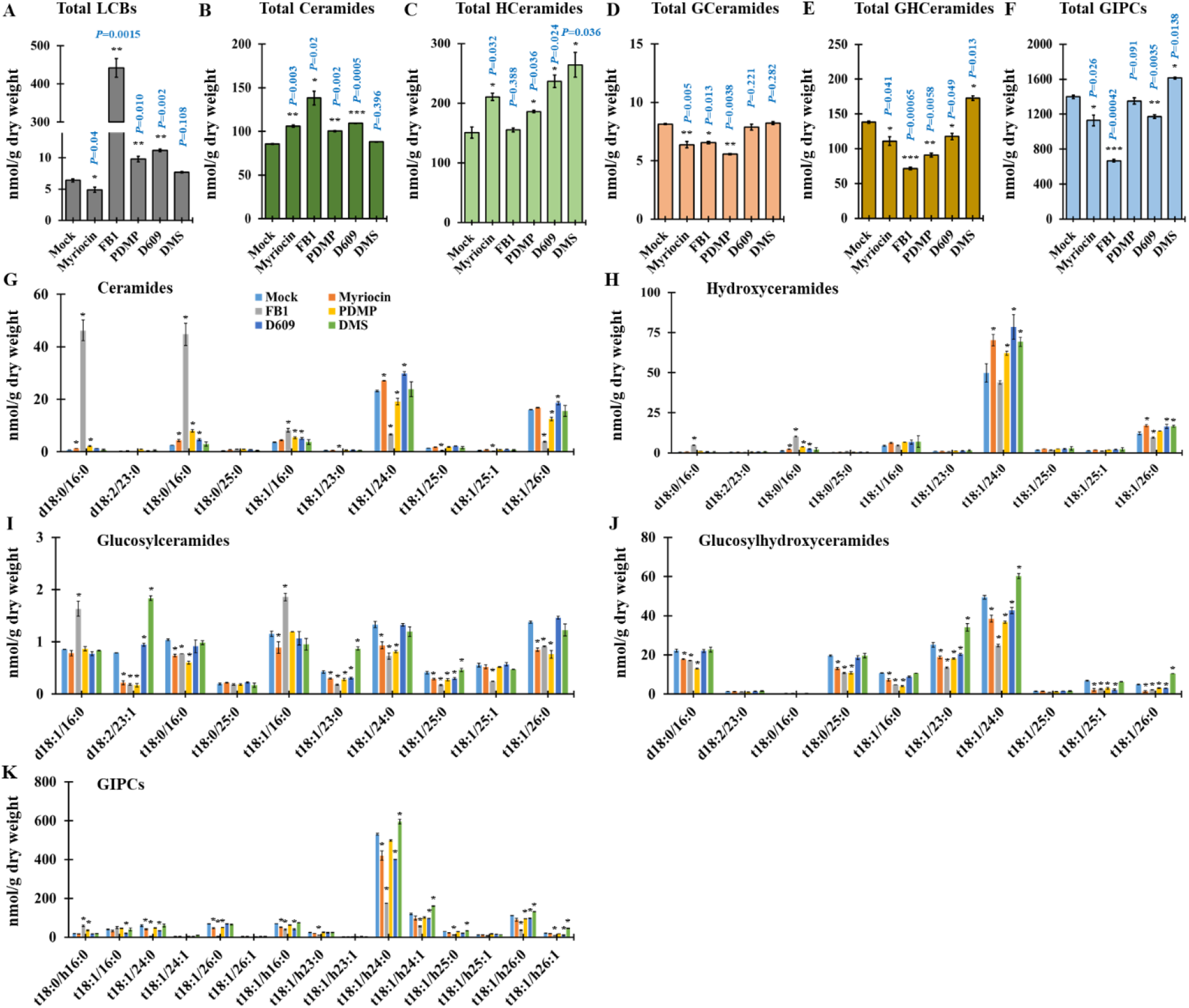
SL inhibitor treatment alters SL content. (**A-F**) Measurement of sphingolipids from wild-type plants after SL inhibitor treatment (24 h) included total LCBs (**A**), total ceramides (**B**), total hydroxyceramides (**C**), total GlcCer (**D**), total GlcHCer (**E**) and total GIPC (**F**). (**G-K**) Sphingolipids species characterized by LCB (d18:0, d18:1, d18:2, t18:0 and t18:1) and fatty acid (FA) (16:0–26:1) from wild type plants after SL inhibitor treatment (24 h) included ceramides (**G**), hydroxyceramides (**H**), GlcCer (**I**), GlcHCer (**J**) and GIPC (**K**). Measurements are the average of four independent biological experiments (n = 200). Data are means ± s.d. Statistical significances were done by two-tailed Student’s t-test; *\*P* < 0.05, *\*\*P* < 0.01, *\*\*\*P* < 0.001.

It was reported that GPI-anchored PdBG2 and PDCB1 proteins are mislocalized in the presence of fenpropimorph (Fen), a potential inhibitor of sterol metabolism that affects lipid raft organization (Grison et al., 2015). Thus, we investigated the subcellular localization of GPI-anchored PD proteins (PdBG2 and PDCB1) after SL inhibitor treatment in *Nicotiana benthamiana* leaves. We constructed several PD reporters for this study (**Fig. 3A**). Firstly, we observed subcellular localization of GFP:PdBG2 and GFP:PDCB1 with PDLP1:RFP, and we found that sterol depletion due to Fen treatment changed the subcellular localization of GFP:PdBG2 and GFP:PDCB1 but not that of PDLP1:RFP (**Supplemental Fig. S3**). Next, we tested the localization of those PD reporters after SL inhibitor treatment. GPI-anchored PdBG2 and PDCB1 were mislocalized in the presence of myriocin, FB1, PDMP and D609, whereas the localization of PDLP1 was not altered (**Fig. 3B-D**). Interestingly, DMS treatment did not change the subcellular localization of any PD reporters (**Fig. 3Q-S**). The β-1,3 glucanase PdBG2 has been shown to regulate callose accumulation, thus, next we asked whether the subcellular localization alteration of this GPI-anchored PdBG2 protein affected the callose level in the SL inhibitor-treated *N. benthamiana* leaves. Our callose analysis showed that the leaves treated myriocin, FB1, PDMP or D609 significantly increased callose accumulation, by contrast DMS-treated leaves exhibited the attenuation of callose accumulation in comparison with mock treatment (**Supplemental Fig. S4**). In addition, we also generated transgenic Arabidopsis plants overexpressing GFP:PdBG2 and observed its subcellular localization after SL inhibitor treatment. Arabidopsis seedlings subjected to the mock or DMS treatment did not change GFP:PdBG2 subcellular localization; the GFP fluorescence pattern was punctuated at PD-PM. Consistently, as shown in the transient expression of GFP:PdBG2, myriocin-, FB1-, PDMP-, or D609-treated seedlings showed mislocalization of GFP:PdBG2, the punctate PD-PM fluorescence pattern was abolished and fluorescence was strongly detected at cytoplasm (**Fig. 4**). Our results suggest that the alteration of GPI-anchored PdBG2 localization is a possible reason that reduce PD callose level after the treatment of a subset of SL pathway inhbitors, resulting in GlcHCers reduction.

**Figure 3.**
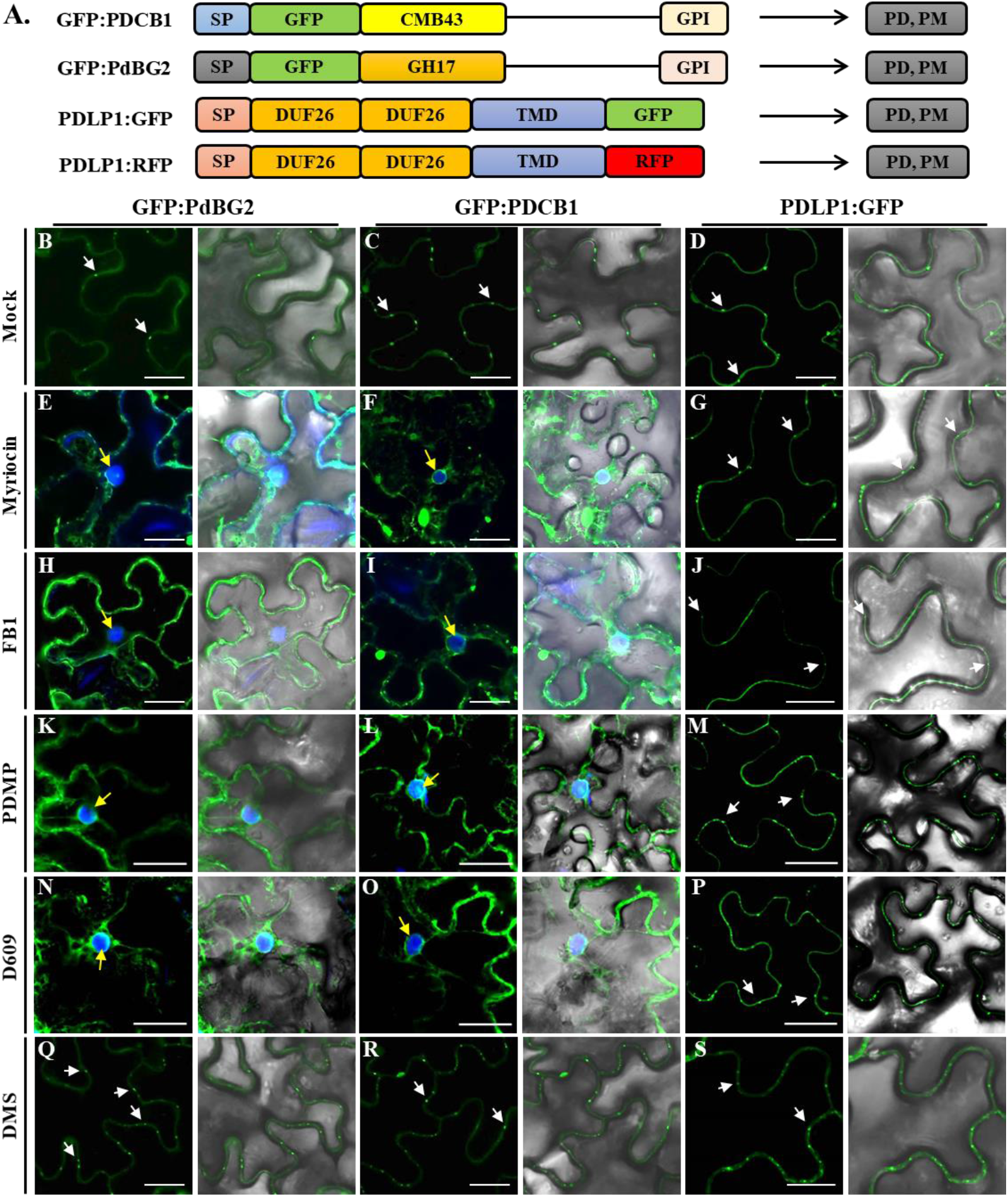
GPI-anchored PD proteins are mislocalized in the presence of SL inhibitors. (**A**) Structural organization of GFP:PDCB1, GFP:PdBG2 and PDLP1:GFP. Chimeric constructs consist of the signal peptides (SP) of PDCB1 and PdBG2, followed by the coding sequence of GFP fused to the callose binding domain (CBM43) for PDCB1, GH17 for PdBG2 and C-terminal GPI anchor signals. For PDLP1, chimeric construct consists of the SP of PDLP1, followed by double DUF26 domain, single transmembrane domain (TMD) and the coding sequence of GFP. (**B-S**) Confocal images of leaf epidermal cells of *N. benthamiana* expressing fluorescent fusion proteins of PdBG2, PDCB1 and PDLP1 after SL inhibitor treatment; (**B, C, D**) mock, (**E, F, G**) 0.1 μM myriocin, (**H, I, J**) 5 μM FB1, (**K, L, M**) 50 μM PDMP, (**N, O, P**) 40 μM D609 and (**Q, R, S**) 30 μM DMS. PD localization (white arrows). DAPI staining was used to determine nuclear localization (yellow arrows). Scale bars = 10 μm.

**Figure 4.**
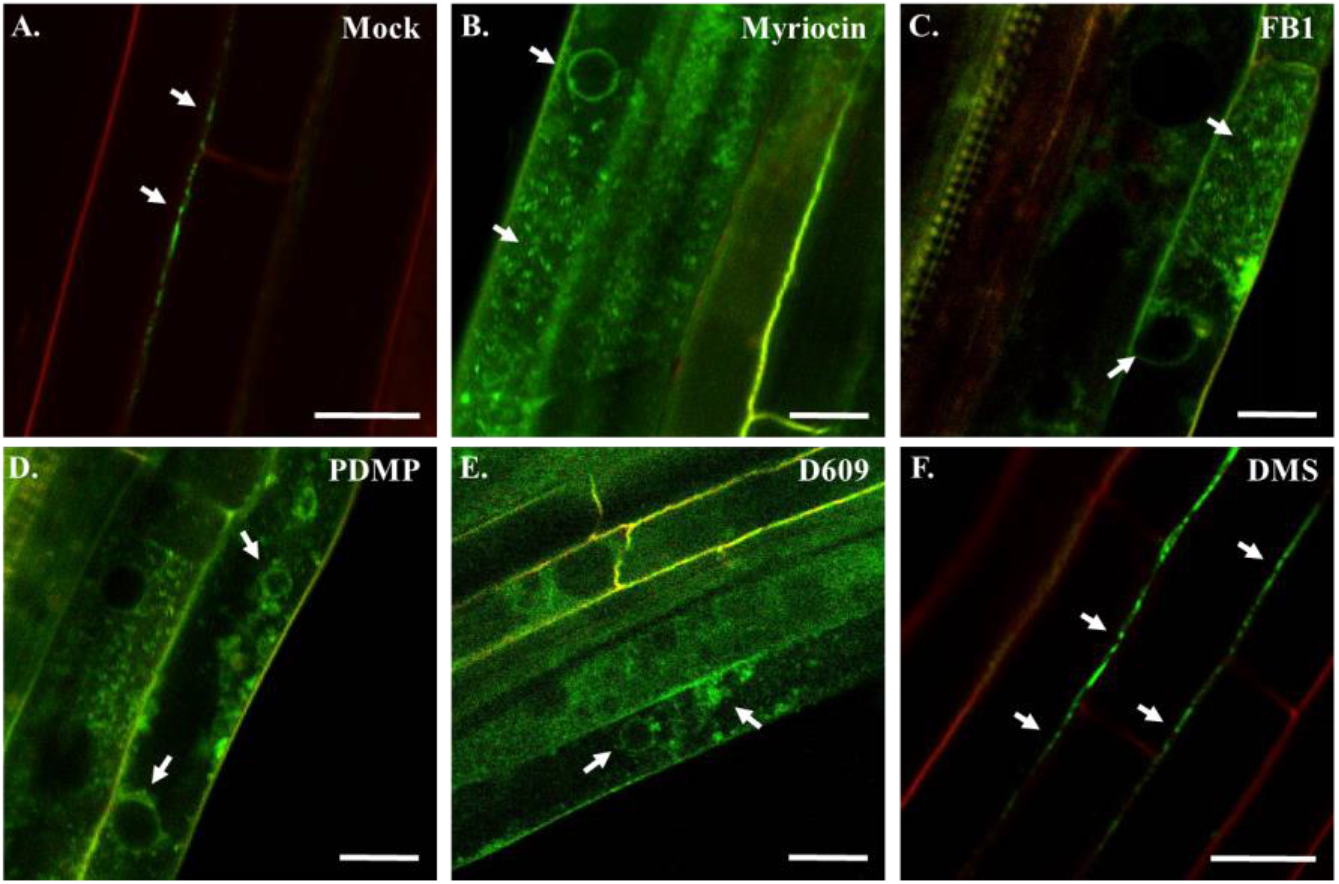
Subcellular localization of PdBG2 after SL inhibitor treatment in Arabidopsis hypocotyls. (**A-F**) Confocal images of Arabidopsis hypocotyls expressing fluorescent fusion protein of PdBG2 after SL inhibitor treatment (24 h); (**A**) mock, (**B**) 0.1 μM myriocin, (**C**) 5 μM FB1, (**D**) 50 μM PDMP, (**E**) 40 μM D609 and (**F**) 30 μM DMS. White arrows indicate GFP:PdBG2 localizations. Red signals showing propidium iodide staining (PI) of the cell wall. Scale bars = 20 μm.

### GlcHCers are associated with PD permeability

In plants, DMS inhibits SPHKs, and PDMP inhibits GCS. In Arabidopsis, SPHK1 was reported to catalyze the formation of sphingosine-1-phosphate (S1P) through the phosphorylation of sphingosine, and GCS is responsible for the formation of GlcCers (Sperling and Heinz, 2003; Silva et al., 2007; Kajiwara et al., 2008; Raffaele et al., 2009; Loizides-Mangold et al., 2012). Based on the SL profiling results after SL inhibitor treatment, GlcHCers were significantly suppressed in the PDMP-treated seedlings condition, however, DMS-treated seedlings firmly showed high GlcHCers in comparison with seedlings subjected to the mock condition. It has been reported that *SPHK1* overexpression plant showed excess callose deposition at Arabidopsis rosette leaves, and it was restricted in the *SPHK1-KD* mutant plant upon fumonisin B1-triggered cell death condition (Qin et al., 2017). Therefore, we employed *sphk1* (*sphk1-2* and *SPHK1-KD*) (**Fig. 5A, C**) and *gcs* mutants in the next experiment to test if the two mutants phenocopy the PD permeability and GlcHCer profiles of the plants treated with DMS and PDMP. The null *gcs* mutant in Arabidopsis showed seedling lethality, but the partial suppression of *GCS* with RNAi resulted in viable and fertile plants (Msanne et al., 2015). We selected another weak mutant allele of the *GCS* gene, *gcs-2,* for which the T-DNA insertion site was predicted to be located in the promoter region, resulting in a slightly reduced transcription level (**Fig. 5B and D**).

**Figure 5.**
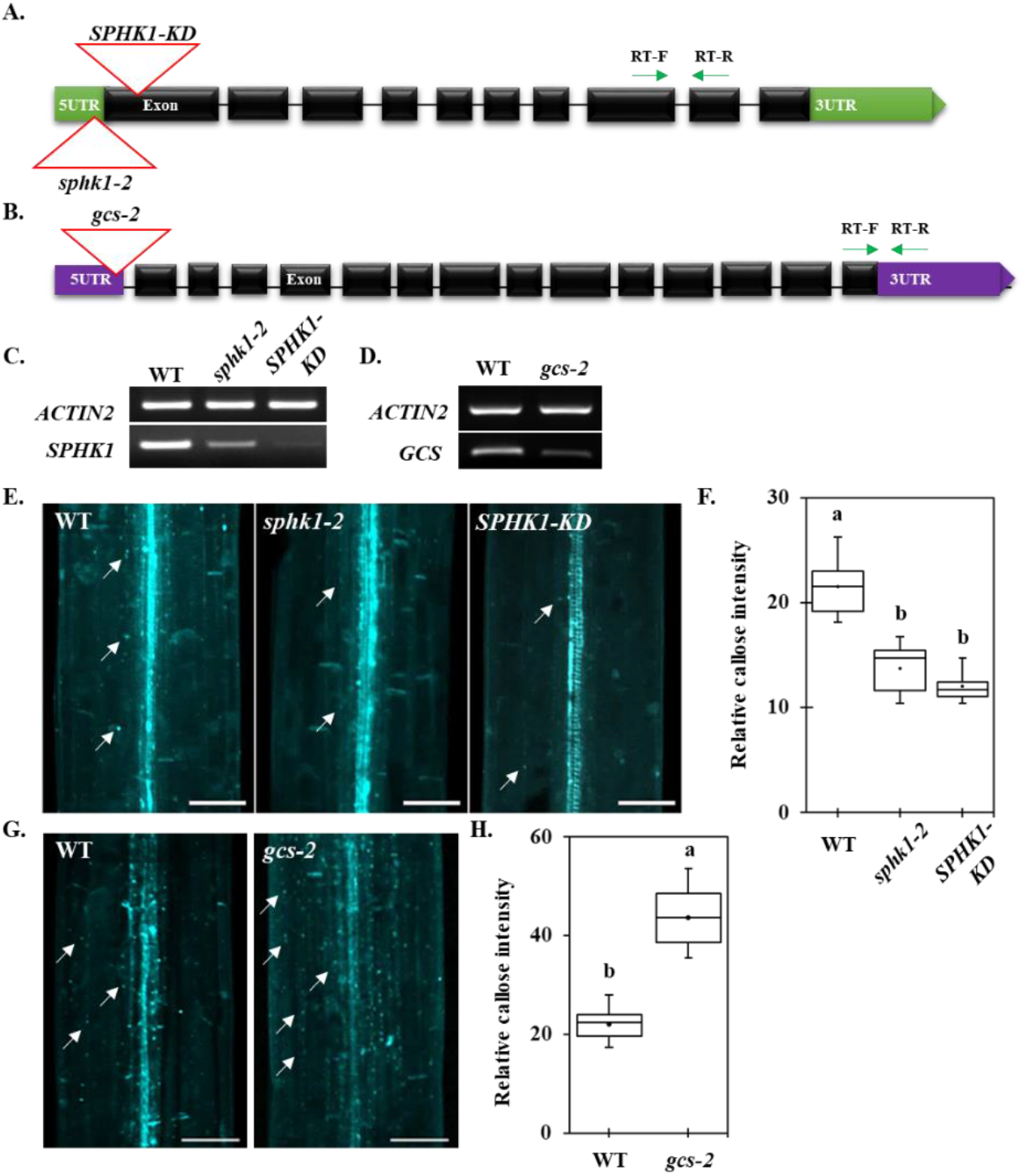
Callose level is altered in the *sphk1-2*, *SPHK1-KD* and *gcs-2* mutants. (**A**) Gene organization of *SPHK1* and its T-DNA insertion alleles. Green arrows indicate RT-primer locations for (**C**). (**B**) Gene organization of *GCS* and its T-DNA insertion allele. Green arrows indicate RT-primer locations for (**D**). (**C**) *SPHK1* transcript expression with alleles containing the T-DNA insertion. (**D**) *GCS* transcript expression with alleles containing the T-DNA insertion. (**E**) Confocal images show aniline blue-stained callose in Arabidopsis etiolated hypocotyls of different genotypes (wild type, *sphk1-2* and *SPHK1-KD*). (**F**) Quantitative data show the relative callose intensity of (**E**). (**G**) Confocal images show aniline blue-stained callose in Arabidopsis etiolated hypocotyls of different genotypes (wild type and *gcs-2*). (**H**) Quantitative data show the relative callose intensity of (**G**). (**E, G**) White arrows show aniline blue-stained callose at PD. Scale bars = 80 μm. (**F, H**) Statistical significances were done by One-Way ANOVA with Tuckey-Kramer test (*n* = 15, 3 independent biological experiments).

To confirm these hypotheses, we tested PD permeability in the *sphk1-2*, *SPHK1-KD* and *gcs-2* mutants by performing HPTS and aniline blue staining at Arabidopsis hypocotyls. The *SPHK1-KD* and *sphk1-2* mutants displayed enhanced PD permeability, depicted by the strong HPTS movement and the reduction in callose deposition (**Fig. 5E, F, Supplemental Fig. S5**). Conversely, the *gcs-2* mutant displayed reduced PD permeability, depicted by the retardation of HPTS movement and the over accumulation of callose deposition (**Fig. 5G, H, Supplemental Fig. S5**). We next conducted SL profile analyses for the *sphk1-2*, *SPHK1-KD* and *gcs-2* mutants. The total contents and several molecules of LCBs, hydroxyceramides, GlcCers, GlcHCers and GIPCs were upregulated in *sphk1-2* and *SPHK1-KD* mutants (**Fig. 6**). On the other hand, the *gcs-2* mutant showed had no significant difference in the LCBs level, but the proportion of hydroxyceramides and GIPCs were significantly enriched in comparison with Col-0 plant. Then, we found a greater attenuation in the proportion of numerous molecules and total contents of GlcCers and GlcHCers (**Fig. 6**), indicating that the *gcs-2* plant also impairs GlcHCers production. Our data showed that the callose deposition phenotype and the proportion of GlcHCers at PDMP- or DMS-treated seedlings were phenocopied by *gcs-2* or *sphk1* (*sphk1-2* and *SPHK1-KD*) mutants, respectively. These data indicate that GlcHCers play the critical role in PD gating via the modulation of callose deposition. We also examined if a perturbation in SL pathways disrupt gene expression of *PdBG2.* Myriocin-, FB1-, PDMP-, or D609-treated seedlings did not change the *PdBG2* expression level as compared with mock treatment, whereas upregulation of *PdBG2* transcript level was observed in the DMS-treated seedlings and *sphk1* mutants (**Supplemental Fig. S6**). This suggests that DMS-treatment or *sphk1* mutation might trigger another signaling pathway in addition to GlcHCers-mediated one.

**Figure 6.**
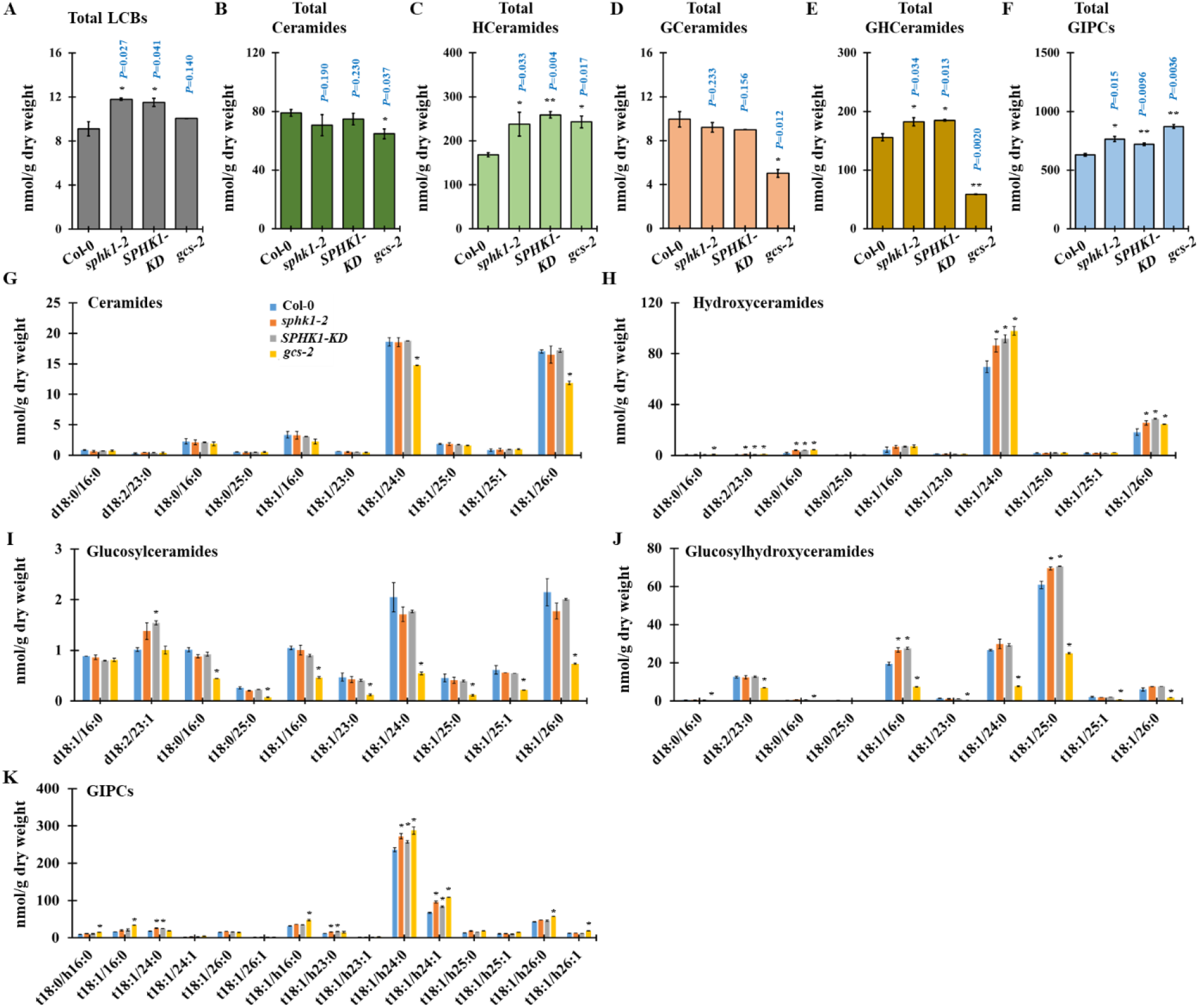
SL homeostasis is altered in the *sphk1-2*, *SPHK1-KD* and *gcs-2* mutants. (**A-F**) Measurement of sphingolipids from *sphk1-2*, *SPHK1-KD* and *gcs-2* mutant plants included total LCBs (A), total ceramides (**B**), total hydroxyceramides (**C**), total GlcCer (**D**), total GlcHCer (**E**) and total GIPC (**F**). (**G-K**) Sphingolipids species characterized by LCB (d18:0, d18:1, d18:2, t18:0 and t18:1) and fatty acid (FA) (16:0–26:1) from *sphk1-2*, *SPHK1-KD* and *gcs-2* mutant plants included ceramides (**G**), hydroxyceramides (**H**), GlcCer (**I**), GlcHCer (**J**) and GIPC (**K**). Measurements are the average of four independent biological experiments (n = 200). Data are means ± s.d. Statistical significances were done by two-tailed Student’s t-test; *\*P* < 0.05, *\*\*P* < 0.01.

### GlcHCers are required for the targeting of GPI-anchored PD proteins

Since myriocin-, FB1-, PDMP-, and D609-suppressed GlcHCers level only disrupted the subcellular localization of the GPI-anchored PD proteins and not that of PDLP1 protein, we hypothesized that the GPI-anchored PdBG2 and PDLP1 proteins might use different cargo machinery in a GlcHCers-enriched vesicle-dependent manner. The secretory pathways for GPI-anchored proteins and non GPI-anchored transmembrane or secretory proteins are distinct in yeast and mammalian cells (Funato and Riezman, 2001; Muniz et al., 2001; Watanabe et al., 2008; Castillon et al., 2009; Rivier et al., 2010; Muniz and Zurzolo, 2014; Paladino et al., 2014; Muniz and Riezman, 2016). During protein secretion in yeast, GPI-anchored proteins are segregated from other proteins and delivered to their final destination by SL-enriched vesicles (Silva et al., 2007; Kajiwara et al., 2008; Loizides-Mangold et al., 2012). So far, there is no clear evidence that GPI-anchored proteins are sorted differently from other transmembrane proteins in plants.

To address the question of whether PdBG2 and PDLP1 proteins are delivered by different pools of secretory vesicles, we used 2-(4-fluorobenzoylamino)-benzoic acid methyl ester (Exo1) to block membrane trafficking between the ER and the Golgi apparatus (Feng et al., 2003; Mishev et al., 2013). First, we examined the subcellular localization of GFP-PdBG2 and PDLP1-GFP after Exo1 treatment in *N. benthamiana* leaves. Cytosolic GFP signals surrounding the nucleus were detected after Exo1 treatment for all proteins (**Supplemental Fig. S7**) and depicted GFP signals seem to be retained in the vesicle compartments. To confirm these results, we next generated fusion protein of well-known vesicle marker involved in the secretory pathway at plant system; VAMP721 protein was selected (Karnahl et al., 2017). We first determined the subcellular localization of VAMP721 protein subjected to mock or Exo1 treatment in *N. benthamiana* leaves. We observed no obvious differences between VAMP721 subcellular localizations in Exo1-treated and control plants (**Supplementary Fig. S8**). To obtain further insight into the mechanisms controlling the secretion of PDLP1 and GPI-anchored PdBG2 proteins, we performed colocalization assays with 3 different combinations; GFP:PdBG2 and VAMP72:RFP, PDLP1:GFP and VAMP721:RFP or GFP:PdBG2 and PDLP1:RFP in the presence of Exo1. Interestingly, the cytosolic GFP signals from PdBG2 and PDLP1 were colocalized with VAMP721:RFP (**Fig. 7A, B**), whereas the cytosolic GFP signal from PdBG2 was not colocalized with RFP signal from PDLP1 (**Fig. 7C**). Our results suggest that there are at least two different vesicles involved in PD protein delivery, and GPI-anchored PdBG2 requires lipid raft-enriched vesicle which is attributed in the GlcHCers composition. (**Supplemental Fig. S9**).

**Figure 7.**
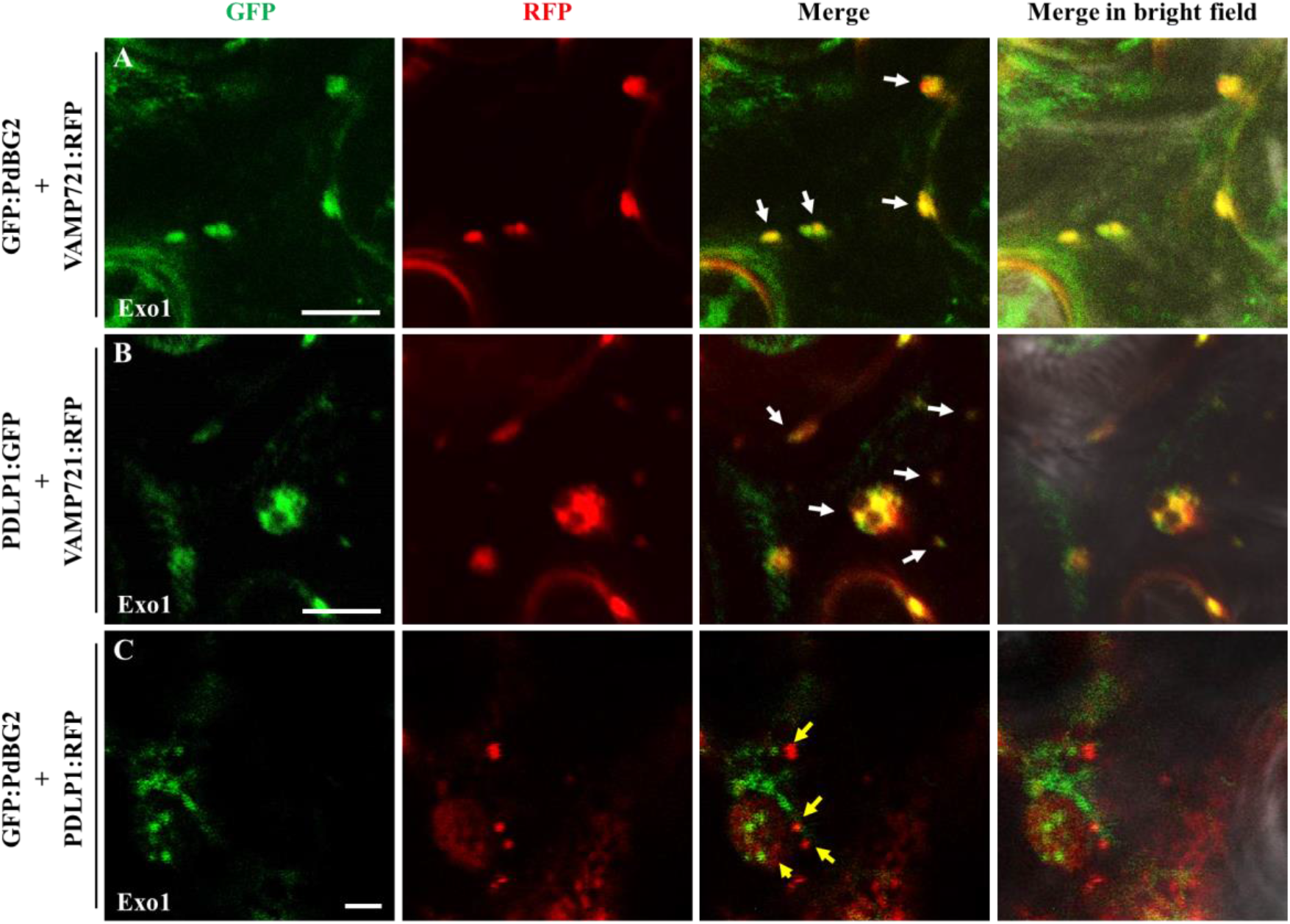
PdBG2 and PDLP1 proteins are secreted with different cargo machinery. (**A**) Confocal images show co-localization between GFP:PdBG2 and VAMP721:RFP after Exo1 (10 μM) treatment (12 h). Scale = 10 µm. (**B**) Confocal images show co-localization between PDLP1:GFP and VAMP721:RFP after Exo1 (10 μM) treatment (12 h). (**A, B**) White arrows indicate GFP and RFP signals were perfectly merged. Scale = 10 µm. (C) Confocal images show different localization between GFP:PdBG2 and PDLP1:RFP after Exo1 (10 μM) treatment (12 h). Yellow arrows indicate GFP and RFP signals were not colocalized. Scale = 2 µm. The fusion proteins were transiently expressed in *N. benthamiana* leaf epidermal cells.

## DISCUSSION

### Cellular localization and transcriptional regulation of PdBG2 determine callose deposition upon GlcHCers alteration

β-1,3-glucanases (BGs) are key players of callose degradation event. There are 50 BG-related genes have been identified in Arabidopsis (Doxey et al., 2007; Kumar et al., 2015). The biological functions of some BGs are tightly connected to the functions of the PD; loss-of-function in PD-localized BGs results in the reduction of the PD SEL caused by excessive callose accumulation in the PD (Doxey et al., 2007; Levy et al., 2007; Levy et al., 2007; Benitez-Alfonso et al., 2013). The most prominent and well-studied of Arabidopsis BGs are AtBG_papp, PdBG1 and PdBG2. Among them, AtBG_papp and PdBG2 are GPI-anchored PD proteins which are attributed to the specialized domains called lipid rafts/microdomains at PD-PM (Levy et al., 2007; Grison et al., 2015). Lipid raft is a unique platform comprised by phospholipids, sterol and SLs that highly enriched in the PD-PM. Moreover, it was shown that PdBG2 protein, which attributes with lipid rafts at PD is mislocalized and induced callose deposition upon sterol depletion (Mongrand et al., 2004; Borner et al., 2005; Grison et al., 2015; Iswanto and Kim, 2017), and this subcellular localization alteration was also observed in the *N. benthamiana* leaves transiently expressing GFP:PdBG2 after Fen treatment (**Supplemental Fig. S3**). Thus, our callose deposition data indicate that perturbation in the SLs composition subjected to the GlcHCers level changes is determined by subcellular localization of PdBG2 protein. Although we could not detect any subcellular localization defect of DMS-treated seedlings, probably there is another molecular mechanism contributes in the GlcHCers-mediated callose deposition.

To address this question, first we investigated the transcriptional activity of *PdBG2* after SLs inhibitor treatment and in the *sphk1* (*SPHK1-KD* and *sphk1-2*) as well as *gcs-2* mutants. Intriguingly, RT-PCR experiment showed that myriocin-, FB1-, PDMP-, or D609-treated seedlings did not change the *PdBG2* expression level as compared with mock treatment, whereas *gcs-2* mutant displayed slightly reduction of *PdBG2* expression in comparison with Col-0 plant. Moreover, upregulation of *PdBG2* transcript level was observed in the DMS-treated seedlings and *sphk1* mutants (**Supplemental Fig. S6**). The latter result implies that post-translational cellular activity of PdBG2 protein is not solely responsible for regulating callose turnover upon GlcHCer fluctuation. Second, it is known that SPHK1 is employed to produce sphingosine-1-phosphate (S1P) (**Supplemental Fig. 1**). In plant, the sphingolipid metabolite S1P involves in guard cell signal transduction (Ng et al., 2001; Coursol et al., 2003; Coursol et al., 2005; Worrall et al., 2008; Puli et al., 2016). S1P in guard triggers alteration in nitric oxide (NO), cytoplasmic calcium and cytoplasmic pH (Ng et al., 2001; Puli et al., 2016). Reactive oxygen species and cytoplasmic calcium are well known trigger of callose accumulation, probably through posttranslational regulation of callose synthases (Xu et al., 2017; Wu et al., 2018). Thus, a reduction of S1P by DMS treatment or in *sphk1* mutants may accompany a diminution of callose synthase activity by posttranslational modification.

Interestingly, recent reports showed that PD-located protein 5 (PDLP5), a member of plasmodesmal-receptor like protein (PD-RLP) family found in Arabidopsis specifically binds to phytosphinganine (t18:0). PDLP5 is highly accumulated in the leaf epidermal cells of *sld1 sld2* double mutant where trihydroxy LCB or phytosphinganine t18:0 is elevated. Similarly, PDLP5 overexpression plant and *sld1 sld2* double mutant show decreased PD permeability through callose accumulation (Chen et al., 2012; Wang et al., 2013; Liu et al., 2020; Vu et al., 2020). However, GlcCers proportions are strongly attenuated in the *sld1 sld2* double mutant (Chen et al., 2012), led us to speculate that GlcHCers might also be impaired. Consistent with our results, reduction in the GlcCers and GlcHCers compositions decrease PD permeability due to an enhancement in callose accumulation which is particularly caused by misregulation of PdBG2 protein.

### Secretory machinery of PdBG2 protein is dependent on the GlcHCers level

This study demonstrates that there are different secretory mechanisms for GPI-anchored PdBG2 protein and non-GPI-anchored PDLP1 protein in plants (**Fig. 7**). The PD proteins harboring the GPI anchor in their C-terminal domain such as PdBG2 and PDCB1 undergo lipid remodeling and GPI modification during protein sorting. The SLs are the LCBs, ceramides, hydroxyceramides, GlcCesr, GlcHCers and GIPCs, and along with sterols, the SLs serve as the basic materials for the formation of lipid rafts or microdomains. In this study, we suggest that lipid raft-enriched vesicles are recruited during the sorting of GPI-anchored PD proteins. Moreover, the attenuation of the more complex SLs such as GlcHCers caused by SL inhibitors (myriocin, FB1, PDMP and D609) or the *gcs* mutation impaired the subcellular localization of GPI-anchored PdBG2 protein. This observation indicates that the sorting of GPI-anchored PdBG2 protein is dependent on GlcHCers proportion.

Since GPI-anchored PD protein targeting is dependent on SLs-enriched lipid rafts, the PDs might be hot spots in which GPI-anchored proteins are enriched. Indeed, many PD proteins have been revealed to be GPI-anchored, including several BGs, the PDCB family members, Remorin (REM), PD localized grain setting defect1 (GSD1) and lysin motif domain-containing glycosyl-phosphatidyl-inositol-anchored protein 2 (LYM2) (Raffaele et al., 2009; Faulkner et al., 2013; Perraki et al., 2014). PDCB1, another GPI-anchored protein, physically binds to callose and is localized in the PDs. Plants overexpressing PDCB1 display enhanced callose accumulation and reduced PD permeability (Simpson et al., 2009). Not all GPI-anchored proteins are PD protein. How GPI-anchored PD proteins and GPI-anchored non-PD proteins are differentially sorted remains to be solved. Our study demonstrated that GlcHCer plays key roles in the formation of the lipid rafts that are required parts of secretion vesicles and that are responsible for the PD targeting of GPI-anchored PdBG2 protein.

## CONCLUSIONS

Overall, our study shows the role of SL molecules in the symplasmic continuity framework. One of important finding is that GlcHCers are keys to the intercellular trafficking machinery of GPI-anchored PD proteins for regulating callose deposition and PD permeability. Our results, demonstrate that subcellular localization of GlcHCers-modulated GPI-anchored PD proteins is dependent on lipid raft integrity. The appropriate function of Exo1 to block the secretory cargo transportation from ER to Golgi enabled us to show that there are at least two different cargo machineries between GPI-anchored PD proteins and non-GPI-anchored PD proteins whereby lipid raft-enriched secretion vesicles is required for transportation of GPI-anchored PD proteins, but not for transportation of non-GPI-anchored PD proteins (**Supplemental Fig. S9**).

## MATERIALS AND METHODS

### Plant material and growth conditions

The *Arabidopsis thaliana* ecotype Colombia-0 (Col-0) and the following mutants were used: *SPHK1-KD* (Sail_794_B01) (Worrall et al., 2008; Qin et al., 2017), *sphk1-2* (Salk_042166), and *gcs-2* (SK31705). The following constructs for the PD reporter were previously described: p35S::SP:GFP:PdBG2 (Grison et al., 2015) and (Simpson et al., 2009), p35S::SP:GFP:PDCB1 (Grison et al., 2015) and (Simpson et al., 2009), p35S::PDLP1:GFP (Caillaud et al., 2014). The pSUC2::GFP construct was transformed into a Col-0 background. Seeds were grown on Murashige and Skoog (MS) (Duchefa Biochemie) agar medium plates (pH 5.9 with KOH, left at 4°C for 2 days in darkness) and then grown in 16-hour light/8-hour dark cycles (long day conditions). *N. benthamiana* plants were grown at 24°C in long day conditions. Plants were grown for 4 weeks before *Agrobacterium tumefaciens*-mediated transient transformation was performed. To generate stable expression in Arabidopsis, a floral dipping method (Clough and Bent, 1998) was used.

### Inhibitor treatments

For the PD permeability analyses (callose deposition and HPTS movement) using Arabidopsis hypocotyls, Col-0 seedlings were grown vertically on normal (without additional SL inhibitors) MS plates for 3 days in the darkness. After 3 days in the dark condition, the seedlings were transferred into MS plates containing the following SL inhibitors: 0.1 µM myriocin (Santa Cruz), 5 µM FB1 (Santa Cruz), 50 µM PDMP (Santa Cruz), 100 µM D609 (Santa Cruz) and 30 µM DMS (Santa Cruz); they were left in dark condition for 12 h prior to measure callose deposition and fluorescence signal intensity. For the PD permeability analyses (callose deposition and GFP fluorescence intensity) using Arabidopsis root tips, Col-0 and Col-0 overexpressing pSUC2::GFP seedlings were grown vertically on normal MS plates for 7 days in light conditions. After 7 days, the seedlings were transferred into MS containing SL inhibitors for 24 h prior to measure callose deposition and GFP signal intensity. Washout experiments were conducted by replaced 24 h-treated seedlings onto MS agar plates subjected to the mock condition for another 72 h before confocal microscopy observation.

### Transient expression analysis

*N. benthamianas* were grown at 24°C under long day conditions. The plants were grown for 4 weeks before *Agobacterium*-mediated transient transformation was performed. Mature 4-week-old leaves were then transformed with the *Agobacterium* strain GV3101 containing the appropriate plasmids, and signals were observed 48 hours after the transformation. Plasmolysis was performed by incubating the leaf samples in 1 M mannitol solution for 15 minutes. The PM was stained with 5 µM FM4-64 solution (Invitrogen Thermo Fisher Scientific) prior to plasmolysis. For the SL inhibitor treatments, 0.1 µM myriocin, 5 µM FB1, 50 µM PDMP, 100 µM D609 or 20 µM DMS were applied at 36 hours after the *Agobacterium* transformation. For the nuclear staining, samples were stained with 8 µg/ml DAPI (Sigma) solution for 10 minutes.

### PD permeability assessments

HPTS loading assays and aniline blue staining ref. (Han et al., 2014; Kumar et al., 2016) were used to assess the permeability of the PD. GFP fluorescence intensity and callose signal intensity were measured using ImageJ software. For additional information on quantification of callose using aniline blue staining, see (Zavaliev et al., 2011; Zavaliev et al., 2013). Data were analyzed by One-Way ANOVA with Tuckey-Kramer test for statistical significance.

### RNA extraction, reverse transcription and RT-PCR

Total RNA from Arabidopsis seedlings (Col-0, *sphk1-2, SPHK1-KD* and *gcs-2*) were extracted using RNeasy^®^ Plant Mini Kit (QIAGEN), according to the manufacturer’s instructions. Aliquots of RNA (1 µg) from each sample were reverse transcribed to cDNA using the QuantiTect^®^ Reverse Transcription Kit (QIAGEN), according to the manufacturer’s instructions. The first strand cDNA was used as a template for quantitative PCR amplification, RT-PCR analyses were conducted using the gene-specific primer pairs under the following conditions: 95°C for 10 minutes followed by 40 cycles of 95°C for 10 seconds, 55°C for 30 seconds, and 72°C for 30 seconds. The expression levels of the different genes were normalized to that of *ACTIN2*. All experiments were repeated independently three times.

### Confocal Microscopy

Confocal fluorescence microscopy was performed with an OLYMPUS FV1000-LDPSU inverted confocal microscope using 20X/0.8 oil-immersion objective or 40X/1.3 oil-immersion objective. GFP was excited with a laser using 488 nanometer beam splitter. RFP and FM4-64 were excited with a laser using 543 nanometer beam splitter. Aniline blue and DAPI were excited with a laser using 405 nanometer beam splitter. Signal intensities from GFP (pSUC2:GFP) and aniline blue (callose detection) were quantified with ImageJ software for statistical data.

### Sphingolipid extraction and sphingolipid analysis

Plant samples preparation for SL inhibitor treatments; seeds (Col-0) were grown on normal MS medium for 14 days. After 10-day-old, seedlings were transferred into MS liquid medium containing mock (DMSO), 0.5 µM Myriocin, 5 µM FB1, 50 µM PDMP, 60 µM D609 and 50 µM DMS and were grown for 24 hours. After 24 hours in SL inhibitor treatments, seedlings were immediately frozen in liquid nitrogen and ground to a fine powder (*n* =250, 4 independent biological experiments). Plant samples preparation for mutant backgrounds (Col-0, *sphk1-1, sphk1-2* and *gcs-2*); seeds were grown on normal MS medium for 14-day-old. Then, 14-day-old seedlings were frozen in liquid nitrogen and ground to a fine powder (*n* = 250, 4 independent biological experiments). For plant sphingolipid analysis, the total lipids were extracted from 3 mg of lyophilized Arabidopsis seedlings using the combined upper phase (220 μL) and lower phase (110 μL) of methyl-*tert*-butyl ether (MTBE)/methanol/water (100:30:35, *v/v/v*) described previously (PMID: 23743007). Extracts were reconstituted in 100 μL chloroform/methanol (1:9, *v/v*). Sphingolipids profiling was performed using a Nexera2 LC system (Shimadzu Corporation, Kyoto, Japan) connected to a triple quadrupole mass spectrometer (LC-MS 8040; Shimadzu, Kyoto, Japan) with reversed phase Kinetex C18 column (100 × 2.1 mm, 2.6 μm, Phenomenex, Torrance, CA, USA) for chromatographic separations of lipids. Mobile phase A consisted of water/methanol (1:9, *v/v*) containing 10 mM ammonium acetate, and mobile phase B consisted of isopropanol/methanol (5:5, *v/v*) containing 10 mM ammonium acetate. To achieve chromatographic separation, a gradient elution program was optimized as follows: 0 min, 30% B; 0-15 min, 95% B; 15-20 min, 95% B; 20-25 min, 30% B. The flow rate was set a 200 μL min^−1^. 5 μL sample volumes were injected for each run. To achieve sphingolipid quantifications, the calculated ratio of analyte and internal standard is multiplied by the concentration of the internal standard to obtain the concentration for each lipid species (PMID: 30835753, 23064709, 28251853, 19517478). We performed quantitative analysis of sphingolipids using one-point calibrations of each target sphingolipid species (dihydrosphingosine d17:0, non-hydroxy phytoceramide 26:0 (t18:0/8:0), 2-hydroxy phytoceramide 24:0 (t18:0/h6:0), and glucosyl-ceramide 30:1 (d18:1/12:0).

## Accession numbers

Accession numbers for the genes characterized in this work are as follows: *PdBG2* (AT2G01630), *PDCB1* (AT5G61130), *PDLP1* (AT5G43980), *SPHK1* (AT4G21540) and *GCS* (AT2G19880).

## Supplemental data

The following supplemental materials are available.

**Supplemental Figure S1.** SL biosynthesis pathways, including their potential inhibitors.

**Supplemental Figure S2.** HPTS movement analysis in the etiolated Arabidopsis hypocotyls.

**Supplemental Figure S3.** GPI-anchored PdBG2 and PDCB1 proteins are mislocalized after Fenpropimorph treatment.

**Supplemental Figure S4.** SL inhibitor treatment alters callose level in *N. benthamiana* leaves.

**Supplemental Figure S5.** HPTS movement analysis in the *sphk1-2*, *SPHK1-KD* and *gcs-2* mutants.

**Supplemental Figure S6.** Transcriptomic analysis of *PdBG2* in the alteration of SL compositions.

**Supplemental Figure S7.** Exo1 inhibits secretory machinery of PdBG2 and PDLP1 proteins.

**Supplemental Figure S8.** Exo1 treatment does not change the cellular localization of VAMP721.

**Supplemental Figure S9.** Schematic model of the role of lipid rafts in the regulation of the PD.

**Supplemental Table S1.** Summary of statistical tests.

**Supplemental Table S2.** The retention times and SRM transitions of the identified *Arabidopsis* sphingolipids.

**Supplemental Table S3.** Distribution of sphingolipids from various tissues of *Arabidopsis*.

**Supplemental Table S4.** Primers used in this study.

## ACKNOWLEDGMENTS

This work was supported by the National Research Foundation of Korea (Grant NRF-2018R1A2A1A05077295 to JYK) and by a grant from the Next-Generation BioGreen 21 Program (SSAC, Grant PJ01137901 to JYK), and the Program for New Plant Breeding Techniques (NBT, Grant PJ01478401), Rural Development Administration (RDA), Republic of Korea.

