## Supplemental Data for "Glucosylhydroxyceramides Modulate Secretion Machinery of a Subset of Plasmodesmata Proteins and a Change in the Callose Accumulation"

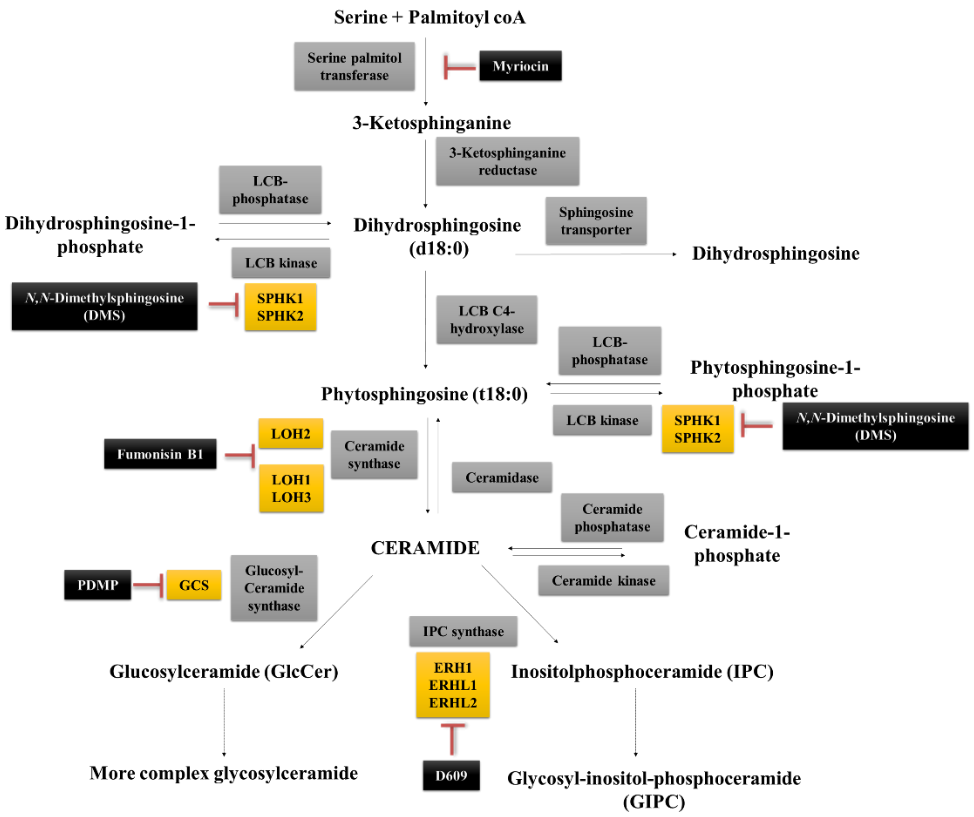


**Supplemental Figure S1. SL biosynthesis pathways, including their potential inhibitors.** Sphingolipid metabolism is started when serine palmitol transferase (SPT) is activated, converting serine and palmitoyl coA to 3-ketosphinganine. This conversion is inhibited by myriocin. 3-Ketosphinganine is further processed to form ceramides in a process facilitated by ceramide synthases. Fumonisin B1 (FB1) from *Fusarium moniliforme* is utilized to attenuate the activity of ceramide synthases. Ceramides need long chain bases (LCBs) as substrates, and dihydrosphingosine (d18:0) and phytosphingosine (t18:0) are the predominant LCBs in plants. These two LCBs are then changed into two different forms of sphingosine-1-phosphate. The formation of sphingosine-1-phosphate is catalyzed by LCB kinases known as SPHINGOSINE KINASEs (SPHKs). *N*, *N*-Dimethyl sphingosine (DMS) appears to be potential inhibitor of SPHK1 activity, decreasing sphingosine-1-phosphate contents. Moreover, ceramide is the precursor for more complex sphingolipid formation, which can be divided into two downstream pathways, namely, the formation of glucosylceramides (GlcCers), which is catalyzed by GLUCOSYL CERAMIDE SYNTHASE (GCS), and the formation of inositol-phospho-ceramides (IPCs), which is catalyzed by IPC SYNTHASE (IPCS). PDMP inhibits the activity of GCS and reduces the level of GlcCers, whereas D609 is known as a phosphatidylcholine-specific phospholipase C (PLC) inhibitor and a sphingomyelin synthase inhibitor in mammalian cells, resulting in the reduction of the level of sphingomyelin.

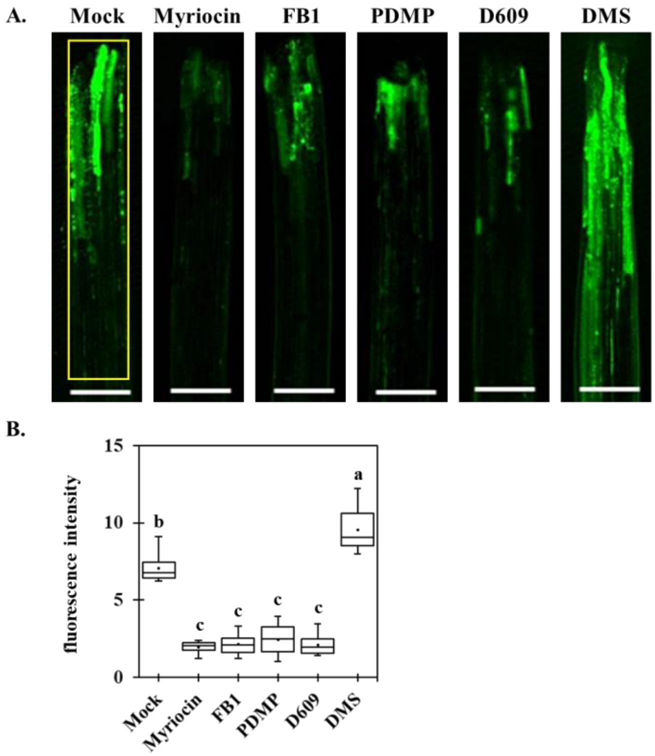


**Supplemental Figure S2. HPTS movement analysis in the etiolated Arabidopsis hypocotyls.**

**(A)** HPTS movement assay was conducted after 12 hours of SL inhibitors treatment (mock, myriocin, FB1, PDMP, D609 and DMS) and the signals were imaged by using excitation at 488 nm. Scale bars = 150 µm.

**(B)** HPTS fluorescence intensity quantification of Arabidopsis hypocotyls **(A)** from each SL inhibitors treatment. Statistical significances were done by One-Way ANOVA with Tuckey-Kramer test (*n* = 30, 3 independent biological experiments; the black dots and lines in boxes indicate the mean and median, respectively).

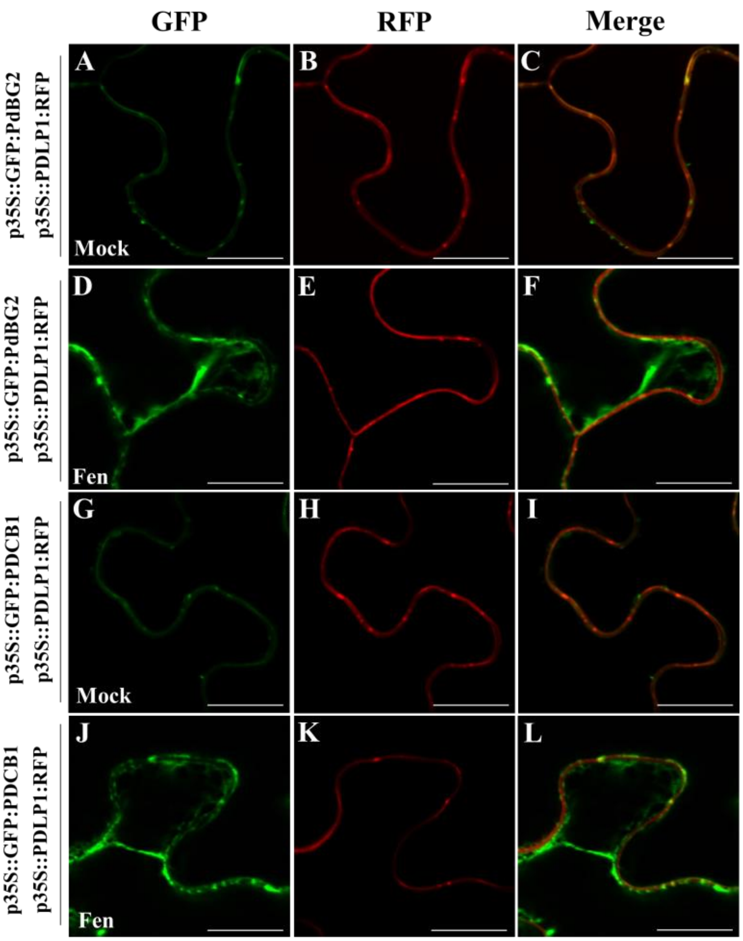


**Supplemental Figure S3. GPI-anchored PdBG2 and PDCB1 proteins are mislocalized after fenpropimorph treatment.**

(**A-C**) Confocal images show co-localization between GFP:PdBG2 (**A**) and PDLP1:RFP (**B**) in *N. benthamiana* leaf epidermal cells in mock condition. White arrows (**C**) show the punctate spots from PdBG2 and PDLP1 are co-localized at PD. Scale bars = 10 µm.

(**D-F**) Confocal images of leaf epidermal cells of *N. benthamiana* expressing fluorescent fusion proteins of GFP:PdBG2 (**D**) and PDLP1:RFP (**F**) after fenpropimorph (fen) treatment (40 µM). The GFP signal is altered (**D**) and not co-localized with RFP signal (**F**). Scale bars = 10 µm.

(**G-I**) Confocal images show co-localization between GFP:PDCB1 (**G**) and PDLP1:RFP (**H**) in *N. benthamiana* leaf epidermal cells in mock condition. White arrows (**I**) show the punctate spots from PDCB1 and PDLP1 are co-localized at PD. Scale bars = 10 µm.

(**J-L**) Confocal images of leaf epidermal cells of *N. benthamiana* expressing fluorescent fusion proteins of GFP:PDCB1 (**J**) and PDLP1:RFP (**K**) after fenpropimorph (fen) treatment (40 µM). The GFP signal is altered (**J**) and not co-localized with RFP signal (**L**). Scale bars = 10 µm.

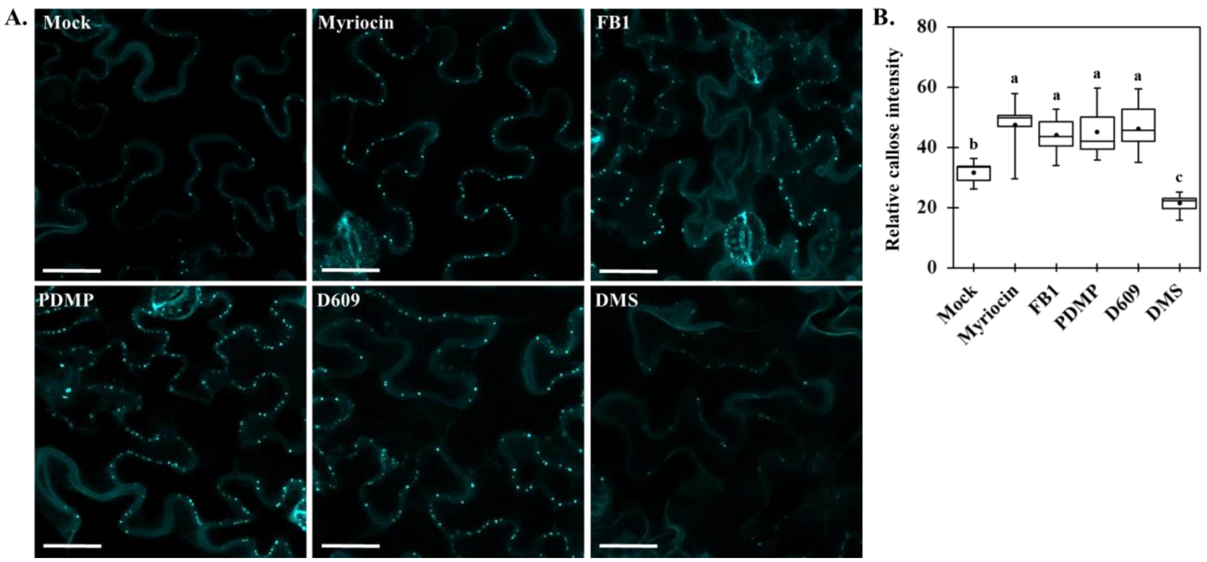


**Supplemental Figure S4. SL inhibitors treatment alters callose level in *N. benthamiana* leaves.** (**A**) Confocal images show aniline blue-stained callose in *N. benthamiana* epidermal cells after SL inhibitors treatment; mock, 0.1 µM myriocin, 5 µM FB1, 50 µM PDMP, 100 µM D609 and 30 µM DMS. Scale bars = 150 µm. (**B**) Quantitative data show the relative callose intensity of (**A**). Statistical significances were done by One-Way ANOVA with Tuckey-Kramer test (*n* = 9, 3 independent biological experiments).

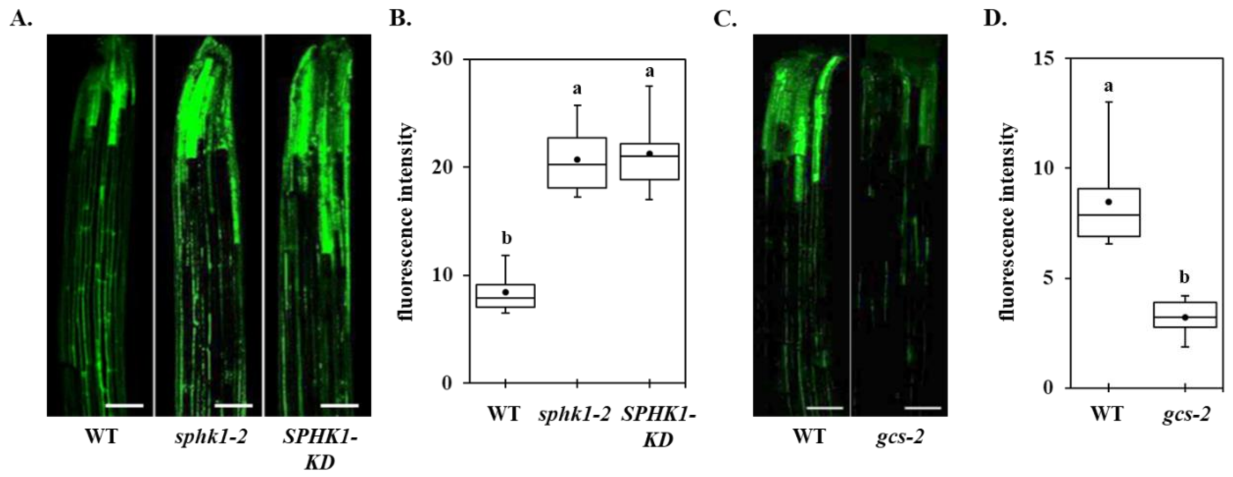


**Supplemental Figure S5. HPTS movement analysis in the *sphk1-2*, *SPHK1-KD* and *gcs-2* mutants.**

(**A**) Confocal images show HPTS movement of wild type, *sphk1-2* and *SPHK1-KD* plants. The signals were imaged by using excitation at 488 nm. Scale bars = 50 µm.

(**B**) Quantification of HPTS fluorescence intensity (**A**). Statistical significances were done by One-Way-ANOVA with Tukey-Kramer test. (*n* = 15, three replicate experiments; the black dots and lines in boxes indicate the mean and median, respectively).

(**C**) Confocal images show HPTS movement of wild type, *gcs-2* plants. Scale bars = 50 µm.

(**D**) Quantification of HPTS fluorescence intensity (**C**). Statistical significances were done by One-Way-ANOVA with Tukey-Kramer test. (*n* = 15, three replicate experiments; the black dots and lines in boxes indicate the mean and median, respectively).

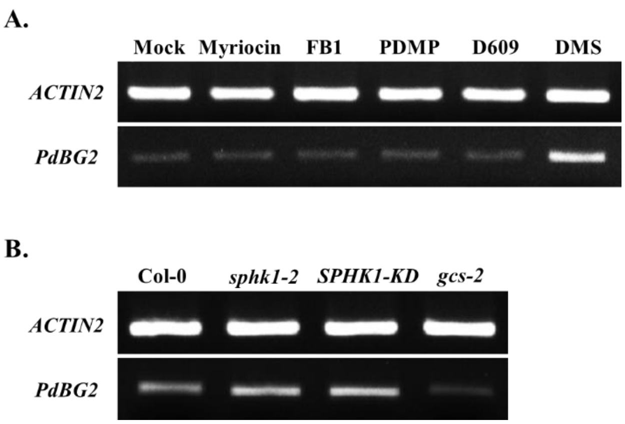


**Supplemental Figure S6. Transcriptomic analysis of *PdBG2* in the alteration of SL compositions.**
(**A**) The transcript levels of *PdBG2* gene in the Arabidopsis seedlings (Col-0) after SL inhibitor treatment.
(**B**) The transcript levels of *PdBG2* gene in *sphk1-2*, *SPHK1-KD* and *gcs-2* mutants.

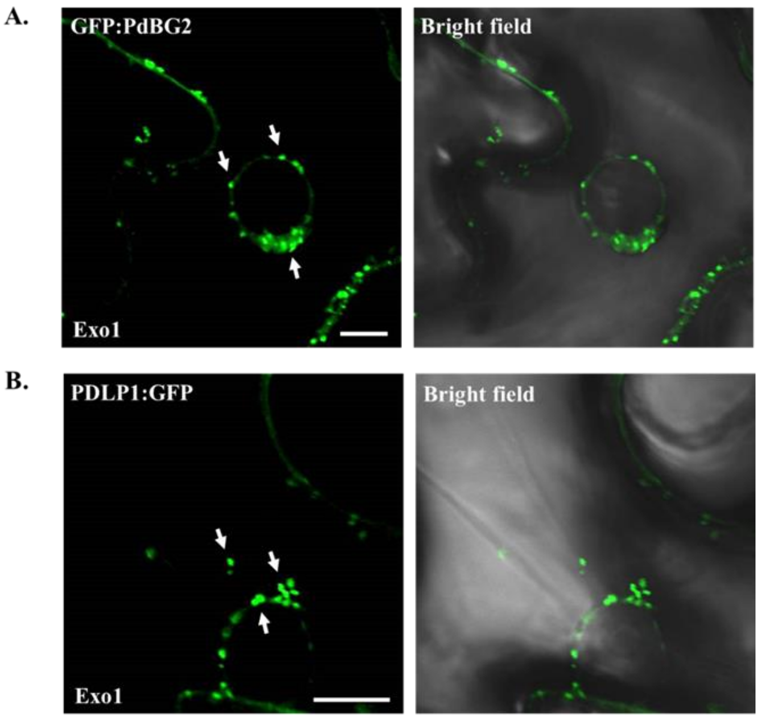


**Supplemental Figure S7. Exo1 inhibits secretory machinery of PdBG2 and PDLP1 proteins.** (**A**) GFP:PdBG2, (**B**) PDLP1:GFP. Confocal images of leaf epidermal cells of *N. benthamiana* transiently expressing GFP fusion proteins after Exo1 treatment (10 μm). White arrows show GFP fusion proteins retained on the cytoplasm. Scale bar = 10 μm.

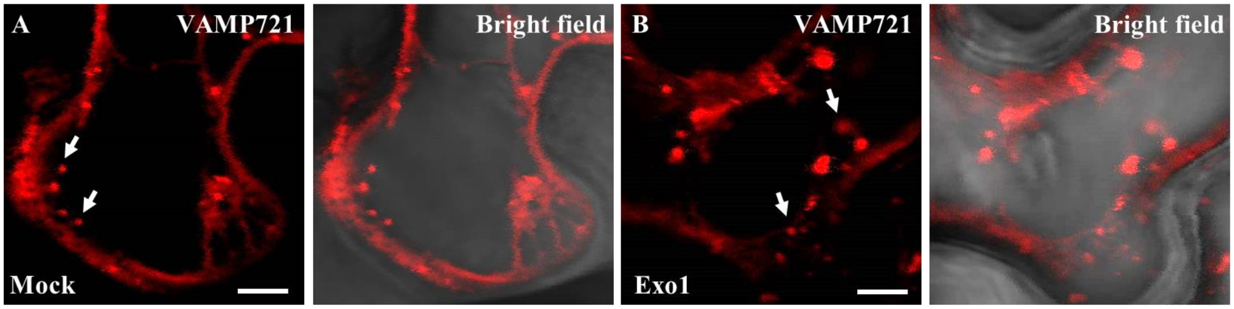


**Supplemental Figure S8. Exo1 treatment does not change the cellular localization of VAMP721.**
(**A, B**) Confocal images show subcellular localization of VAMP721:RFP in the absence (**A**) and presence (**B**) of Exo1 (10 μM). VAMP721:RFP was transiently expressed in *N. benthamiana* leaf epidermal cells. Scale = 5 μm.

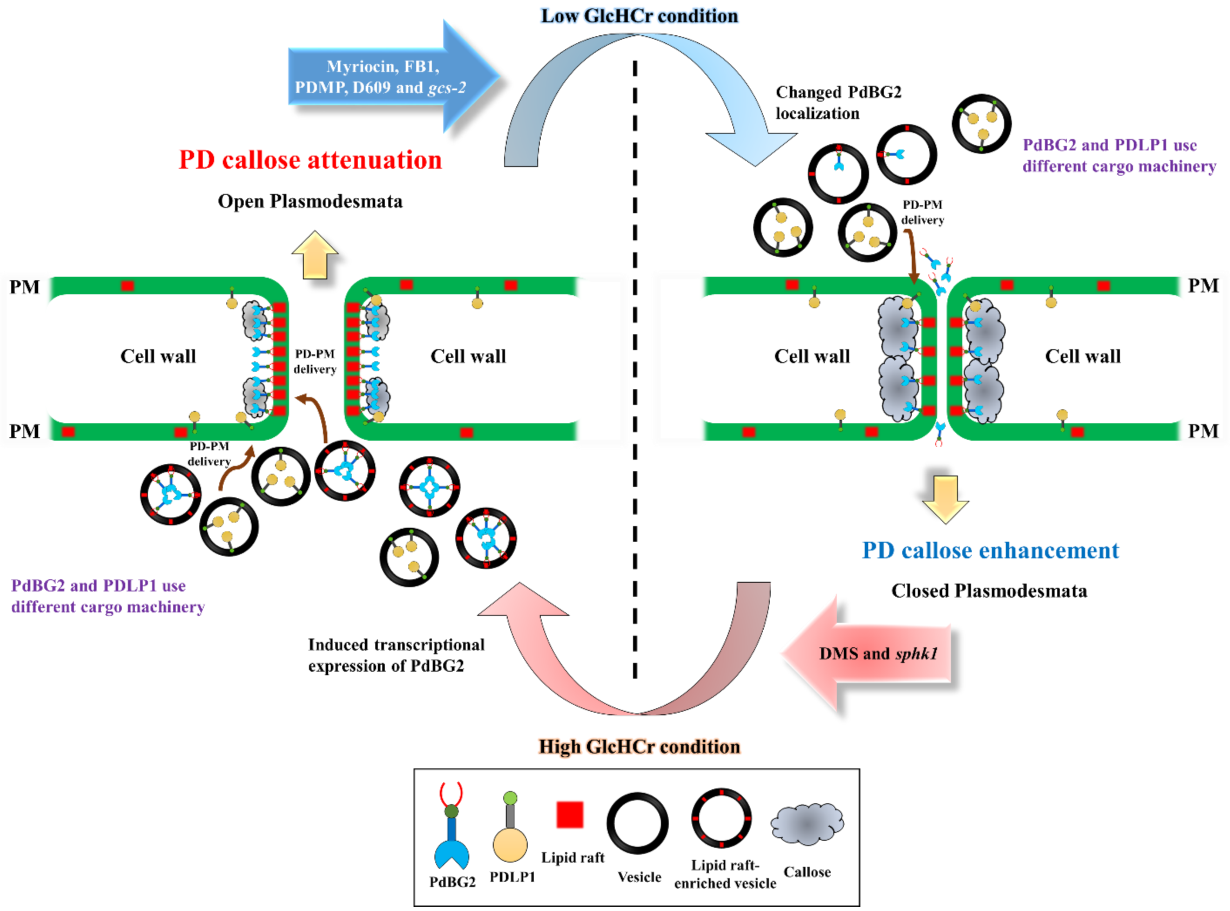


**Supplemental Figure S9. Schematic model of the role of lipid rafts in the regulation of the PD.** PD-PM is enriched in saturated lipid moieties, and the reduction of GlcHCer at *gcs-2*, myriocin, FB1, PDMP and D609 conditions led to cellular localization alteration of GPI-anchored PdBG2 (not for PDLP1) thus induces callose level at PD. Meanwhile, high GlcHCer condition at *sphk1* and DMS treatment do not change either PdBG2 or PDLP1 cellular localization, but transcriptional expression of PdBG2 is upregulated thus might induces callose degradation activity and open PD. Lipid raft-enriched vesicles are the designated platforms that enable GPI-anchored PdBG2 to reach their destination in the PD. Moreover, the attenuation of the more complex SL such as GlcHCer affect SL homeostasis, triggering disassembly of the lipid rafts and alteration in the distribution of GPI-anchored PdBG2.

**Supplemental Table S2.** The retention times and SRM transitions of the identified Arabidopsis sphingolipids

| **No.** | **Class** | **Species** | **Adduct ion** | **$Q_1$ (*m/z*)** | **$Q_3$ (*m/z*)** | **CE (eV)** | **Retention time (min)** |
|---|---|---|---|---|---|---|---|
| 1 | LCB* | d17:0 | $[M+H]^+$ | 288.3 | 270.3 | 12 | 1.5 |
| 2 | CER* | CER t18:0/8:0 | $[M+H]^+$ | 444.6 | 282.3 | 32 | 2.1 |
| 3 | HCER* | HCER t18:0/h6:0 | $[M+H]^+$ | 432.6 | 282.3 | 32 | 1.7 |
| 4 | GCER* | GCER d18:1/12:0 | $[M+H]^+$ | 644.6 | 264.3 | 35 | 2.9 |
| 5 | GIPC* | $GM_1$ d18:1/18:0 | $[M+H]^+$ | 1546.9 | 366.1 | 30 | 4.2 |
| 6 | LCB | d18:0 | $[M+H]^+$ | 302.3 | 284.3 | 12 | 1.6 |
| 7 | LCB | t18:0 | $[M+H]^+$ | 318.3 | 300.3 | 12 | 1.4 |
| 8 | LCB | t18:1 | $[M+H]^+$ | 316.3 | 298.3 | 12 | 1.4 |
| 9 | CER | CER d18:0/16:0 | $[M+H]^+$ | 540.5 | 266.2 | 32 | 7.3 |
| 10 | CER | CER d18:0/24:0 | $[M+H]^+$ | 652.6 | 266.2 | 32 | 12.6 |
| 11 | CER | CER d18:1/22:0 | $[M+H]^+$ | 622.6 | 264.3 | 32 | 10.9 |
| 12 | CER | CER d18:1/24:0 | $[M+H]^+$ | 650.6 | 264.3 | 32 | 12.2 |
| 13 | CER | CER d18:2/23:0 | $[M+H]^+$ | 634.6 | 262.3 | 32 | 7.4 |
| 14 | CER | CER d18:2/25:0 | $[M+H]^+$ | 662.6 | 262.3 | 32 | 8.0 |
| 15 | CER | CER t18:0/16:0 | $[M+H]^+$ | 556.5 | 282.3 | 32 | 6.1 |
| 16 | CER | CER t18:0/18:2 | $[M+H]^+$ | 580.5 | 282.3 | 32 | 7.4 |
| 17 | CER | CER t18:0/22:0 | $[M+H]^+$ | 640.5 | 282.3 | 32 | 10.2 |
| 18 | CER | CER t18:0/23:0 | $[M+H]^+$ | 654.5 | 282.3 | 32 | 10.8 |
| 19 | CER | CER t18:0/24:0 | $[M+H]^+$ | 668.6 | 282.3 | 32 | 11.5 |
| 20 | CER | CER t18:0/24:1 | $[M+H]^+$ | 666.6 | 282.3 | 32 | 10.2 |
| 21 | CER | CER t18:0/25:0 | $[M+H]^+$ | 682.6 | 282.3 | 32 | 12.1 |
| 22 | CER | CER t18:0/25:1 | $[M+H]^+$ | 680.6 | 282.3 | 32 | 10.9 |
| 23 | CER | CER t18:0/26:0 | $[M+H]^+$ | 696.6 | 282.3 | 32 | 12.7 |
| 24 | CER | CER t18:1/16:0 | $[M+H]^+$ | 554.5 | 280.3 | 32 | 5.5 |
| 25 | CER | CER t18:1/22:0 | $[M+H]^+$ | 638.5 | 280.3 | 32 | 9.5 |
| 26 | CER | CER t18:1/23:0 | $[M+H]^+$ | 652.5 | 280.3 | 32 | 10.2 |
| 27 | CER | CER t18:1/24:0 | $[M+H]^+$ | 666.6 | 280.3 | 32 | 10.8 |
| 28 | CER | CER t18:1/24:1 | $[M+H]^+$ | 664.6 | 280.3 | 32 | 9.6 |
| 29 | CER | CER t18:1/25:0 | $[M+H]^+$ | 680.6 | 280.3 | 32 | 11.5 |
| 30 | CER | CER t18:1/25:1 | $[M+H]^+$ | 678.6 | 280.3 | 32 | 10.3 |

| | | | | | | | |
|---|---|---|---|---|---|---|---|
| 31 | CER | CER t18:1/26:0 | $[M+H]^+$ | 694.6 | 280.3 | 32 | 12.1 |
| 32 | HCER | HCER d18:0/h16:0 | $[M+H]^+$ | 556.5 | 266.2 | 32 | 6.9 |
| 33 | HCER | HCER d18:1/h16:0 | $[M+H]^+$ | 554.5 | 264.3 | 32 | 5.3 |
| 34 | HCER | HCER d18:1/h24:0 | $[M+H]^+$ | 666.6 | 264.3 | 32 | 10.2 |
| 35 | HCER | HCER d18:2/h16:0 | $[M+H]^+$ | 552.5 | 262.3 | 32 | 4.3 |
| 36 | HCER | HCER d18:2/h22:1 | $[M+H]^+$ | 634.6 | 262.3 | 32 | 7.4 |
| 37 | HCER | HCER d18:2/h24:0 | $[M+H]^+$ | 664.6 | 262.3 | 32 | 9.6 |
| 38 | HCER | HCER d18:2/h24:1 | $[M+H]^+$ | 662.6 | 262.3 | 32 | 8.0 |
| 39 | HCER | HCER d18:2/h26:0 | $[M+H]^+$ | 692.7 | 262.3 | 32 | 10.9 |
| 40 | HCER | HCER t18:0/h16:0 | $[M+H]^+$ | 572.5 | 282.3 | 32 | 5.6 |
| 41 | HCER | HCER t18:0/h22:0 | $[M+H]^+$ | 656.5 | 282.3 | 32 | 9.6 |
| 42 | HCER | HCER t18:0/h22:1 | $[M+H]^+$ | 654.6 | 282.3 | 32 | 8.4 |
| 43 | HCER | HCER t18:0/h23:0 | $[M+H]^+$ | 670.5 | 282.3 | 32 | 10.3 |
| 44 | HCER | HCER t18:0/h23:1 | $[M+H]^+$ | 668.5 | 282.3 | 32 | 11.5 |
| 45 | HCER | HCER t18:0/h24:0 | $[M+H]^+$ | 684.6 | 282.3 | 32 | 10.9 |
| 46 | HCER | HCER t18:0/h24:1 | $[M+H]^+$ | 682.6 | 282.3 | 32 | 9.7 |
| 47 | HCER | HCER t18:0/h25:0 | $[M+H]^+$ | 698.6 | 282.3 | 32 | 11.6 |
| 48 | HCER | HCER t18:0/h25:1 | $[M+H]^+$ | 696.6 | 282.3 | 32 | 10.3 |
| 49 | HCER | HCER t18:0/h26:0 | $[M+H]^+$ | 712.6 | 282.3 | 32 | 12.2 |
| 50 | HCER | HCER t18:0/h26:1 | $[M+H]^+$ | 710.6 | 282.3 | 32 | 11.0 |
| 51 | HCER | HCER t18:1/h16:0 | $[M+H]^+$ | 570.5 | 280.3 | 32 | 4.3 |
| 52 | HCER | HCER t18:1/h22:0 | $[M+H]^+$ | 654.5 | 280.3 | 32 | 7.9 |
| 53 | HCER | HCER t18:1/h22:1 | $[M+H]^+$ | 652.6 | 280.3 | 32 | 6.8 |
| 54 | HCER | HCER t18:1/h23:0 | $[M+H]^+$ | 668.5 | 280.3 | 32 | 9.6 |
| 55 | HCER | HCER t18:1/h23:1 | $[M+H]^+$ | 666.5 | 280.3 | 32 | 10.9 |
| 56 | HCER | HCER t18:1/h24:0 | $[M+H]^+$ | 682.6 | 280.3 | 32 | 9.2 |
| 57 | HCER | HCER t18:1/h24:1 | $[M+H]^+$ | 680.6 | 280.3 | 32 | 8.0 |
| 58 | HCER | HCER t18:1/h25:0 | $[M+H]^+$ | 696.6 | 280.3 | 32 | 11.0 |
| 59 | HCER | HCER t18:1/h25:1 | $[M+H]^+$ | 694.6 | 280.3 | 32 | 12.1 |
| 60 | HCER | HCER t18:1/h26:0 | $[M+H]^+$ | 710.6 | 280.3 | 32 | 10.5 |
| 61 | HCER | HCER t18:1/h26:1 | $[M+H]^+$ | 708.6 | 280.3 | 32 | 9.3 |
| 62 | GCER | GCER d18:1/16:0 | $[M+H]^+$ | 700.5 | 264.3 | 35 | 5.6 |
| 63 | GCER | GCER d18:2/23:1 | $[M+H]^+$ | 794.6 | 262.3 | 35 | 5.0 |
| 64 | GCER | GCER t18:0/16:0 | $[M+H]^+$ | 718.5 | 282.3 | 35 | 5.4 |

| | | | | | | | |
|---|---|---|---|---|---|---|---|
| 65 | GCER | GCER t18:0/25:0 | $[M+H]^+$ | 844.5 | 282.3 | 35 | 8.1 |
| 66 | GCER | GCER t18:1/16:0 | $[M+H]^+$ | 716.5 | 280.3 | 35 | 4.6 |
| 67 | GCER | GCER t18:1/23:0 | $[M+H]^+$ | 814.5 | 280.3 | 35 | 6.8 |
| 68 | GCER | GCER t18:1/24:0 | $[M+H]^+$ | 828.5 | 280.3 | 35 | 7.4 |
| 69 | GCER | GCER t18:1/25:0 | $[M+H]^+$ | 842.5 | 280.3 | 35 | 8.0 |
| 70 | GCER | GCER t18:1/25:1 | $[M+H]^+$ | 840.6 | 280.3 | 35 | 7.1 |
| 71 | GCER | GCER t18:1/26:0 | $[M+H]^+$ | 856.5 | 280.3 | 35 | 8.7 |
| 72 | GHCER | GHCER d18:0/h16:0 | $[M+H]^+$ | 718.5 | 266.2 | 35 | 6.0 |
| 73 | GHCER | GHCER d18:1/h16:0 | $[M+H]^+$ | 716.5 | 264.3 | 35 | 5.4 |
| 74 | GHCER | GHCER d18:1/h24:1 | $[M+H]^+$ | 826.6 | 264.3 | 35 | 9.2 |
| 75 | GHCER | GHCER d18:2/h16:0 | $[M+H]^+$ | 714.5 | 262.3 | 35 | 4.2 |
| 76 | GHCER | GHCER t18:0/h16:0 | $[M+H]^+$ | 734.5 | 282.3 | 35 | 4.9 |
| 77 | GHCER | GHCER t18:0/h24:0 | $[M+H]^+$ | 846.5 | 282.3 | 35 | 9.2 |
| 78 | GHCER | GHCER t18:0/h24:1 | $[M+H]^+$ | 844.6 | 282.3 | 35 | 8.0 |
| 79 | GHCER | GHCER t18:1/h16:0 | $[M+H]^+$ | 732.5 | 280.3 | 35 | 4.3 |
| 80 | GHCER | GHCER t18:1/h18:0 | $[M+H]^+$ | 760.5 | 280.3 | 35 | 5.4 |
| 81 | GHCER | GHCER t18:1/h20:0 | $[M+H]^+$ | 788.5 | 280.3 | 35 | 6.6 |
| 82 | GHCER | GHCER t18:1/h22:0 | $[M+H]^+$ | 816.5 | 280.3 | 35 | 7.9 |
| 83 | GHCER | GHCER t18:1/h22:1 | $[M+H]^+$ | 814.6 | 280.3 | 35 | 6.8 |
| 84 | GHCER | GHCER t18:1/h23:0 | $[M+H]^+$ | 830.5 | 280.3 | 35 | 8.6 |
| 85 | GHCER | GHCER t18:1/h23:1 | $[M+H]^+$ | 828.6 | 280.3 | 35 | 7.4 |
| 86 | GHCER | GHCER t18:1/h24:0 | $[M+H]^+$ | 844.5 | 280.3 | 35 | 9.2 |
| 87 | GHCER | GHCER t18:1/h24:1 | $[M+H]^+$ | 842.6 | 280.3 | 35 | 8.0 |
| 88 | GHCER | GHCER t18:1/h25:0 | $[M+H]^+$ | 858.5 | 280.3 | 35 | 9.9 |
| 89 | GHCER | GHCER t18:1/h25:1 | $[M+H]^+$ | 856.6 | 280.3 | 35 | 8.7 |
| 90 | GHCER | GHCER t18:1/h26:0 | $[M+H]^+$ | 872.5 | 280.3 | 35 | 10.5 |
| 91 | GHCER | GHCER t18:1/h26:1 | $[M+H]^+$ | 870.6 | 280.3 | 35 | 9.3 |
| 92 | GIPC | GIPC t18:0/h16:0 | $[M+H]^+$ | 1152.6 | 554.5 | 30 | 2.3 |
| 93 | GIPC | GIPC t18:0/h22:0 | $[M+H]^+$ | 1236.7 | 638.6 | 30 | 4.6 |
| 94 | GIPC | GIPC t18:0/h22:1 | $[M+H]^+$ | 1234.7 | 636.6 | 30 | 3.2 |
| 95 | GIPC | GIPC t18:0/h23:0 | $[M+H]^+$ | 1250.7 | 652.6 | 30 | 5.2 |
| 96 | GIPC | GIPC t18:0/h23:1 | $[M+H]^+$ | 1248.7 | 650.6 | 30 | 4.2 |
| 97 | GIPC | GIPC t18:0/h24:0 | $[M+H]^+$ | 1264.8 | 666.7 | 30 | 5.8 |
| 98 | GIPC | GIPC t18:0/h24:1 | $[M+H]^+$ | 1262.8 | 664.7 | 30 | 4.8 |

| | | | | | | | |
|---|---|---|---|---|---|---|---|
| 99 | GIPC | GIPC t18:0/h25:0 | $[M+H]^+$ | 1278.8 | 680.7 | 30 | 6.4 |
| 100 | GIPC | GIPC t18:0/h25:1 | $[M+H]^+$ | 1276.8 | 678.7 | 30 | 5.4 |
| 101 | GIPC | GIPC t18:0/h26:0 | $[M+H]^+$ | 1292.8 | 694.7 | 30 | 7.1 |
| 102 | GIPC | GIPC t18:0/h26:1 | $[M+H]^+$ | 1290.8 | 692.7 | 30 | 6.0 |
| 103 | GIPC | GIPC t18:1/16:0 | $[M+H]^+$ | 1134.6 | 536.6 | 30 | 2.4 |
| 104 | GIPC | GIPC t18:1/22:0 | $[M+H]^+$ | 1218.7 | 620.6 | 30 | 4.5 |
| 105 | GIPC | GIPC t18:1/24:0 | $[M+H]^+$ | 1246.8 | 648.7 | 30 | 5.7 |
| 106 | GIPC | GIPC t18:1/24:1 | $[M+H]^+$ | 1244.7 | 646.6 | 30 | 4.7 |
| 107 | GIPC | GIPC t18:1/26:0 | $[M+H]^+$ | 1274.8 | 676.7 | 30 | 7.0 |
| 108 | GIPC | GIPC t18:1/26:1 | $[M+H]^+$ | 1272.8 | 674.7 | 30 | 5.8 |
| 109 | GIPC | GIPC t18:1/h16:0 | $[M+H]^+$ | 1150.6 | 552.5 | 30 | 2.1 |
| 110 | GIPC | GIPC t18:1/h20:0 | $[M+H]^+$ | 1206.7 | 608.6 | 30 | 3.2 |
| 111 | GIPC | GIPC t18:1/h22:0 | $[M+H]^+$ | 1234.7 | 636.6 | 30 | 4.1 |
| 112 | GIPC | GIPC t18:1/h22:1 | $[M+H]^+$ | 1232.7 | 634.6 | 30 | 3.4 |
| 113 | GIPC | GIPC t18:1/h23:0 | $[M+H]^+$ | 1248.7 | 650.6 | 30 | 4.7 |
| 114 | GIPC | GIPC t18:1/h23:1 | $[M+H]^+$ | 1246.7 | 648.6 | 30 | 3.8 |
| 115 | GIPC | GIPC t18:1/h24:0 | $[M+H]^+$ | 1262.8 | 664.7 | 30 | 5.3 |
| 116 | GIPC | GIPC t18:1/h24:1 | $[M+H]^+$ | 1260.8 | 662.7 | 30 | 4.2 |
| 117 | GIPC | GIPC t18:1/h25:0 | $[M+H]^+$ | 1276.8 | 678.7 | 30 | 5.9 |
| 118 | GIPC | GIPC t18:1/h25:1 | $[M+H]^+$ | 1274.8 | 676.7 | 30 | 4.8 |
| 119 | GIPC | GIPC t18:1/h26:0 | $[M+H]^+$ | 1290.8 | 692.7 | 30 | 6.5 |
| 120 | GIPC | GIPC t18:1/h26:1 | $[M+H]^+$ | 1288.8 | 690.7 | 30 | 5.4 |

*IS (Internal standard)

**Supplemental Table S3.** Distribution of sphingolipids from various tissues of *Arabidopsis*.

| **Reference (PMID)** | Our result | Wu JX[1] (26734030) | Tellier F (24731258) | Luttgearm KD (25794895) | Luttgearm KD (26313010) | Xie LJ (25822663) | Markham JE (17340572) | Markham JE (16772288) | Chao DY (21421810) |
|---|---|---|---|---|---|---|---|---|---|
| Tissue | Whole | Whole | Whole | pollens | leaves & calli | rossette | leaves | leaves | hypocotyls |
| Growth conditions | 9 days | 6 days | 12 days | 5 weeks | 4 weeks | 4 weeks | 5-6 weeks | 5-6 weeks | 5 weeks |
| **Lipid species (nmol/g dry wt)[2]** | | | | | | | | | |
| CERs | 85.5 ± 0.8 | 70 | 60 | 107 ± 18 | 170 | 15 | 16.3 ± 1.3 | 13.7 | 30 |
| HCERs | 150.8 ± 18.8 | 68 | 90 | 345 ± 79 | 220 | 40 | 12.4 ± 0.8 | - | 25 |
| GCERs | 146.5 ± 2.1 | - | 165 | 1377 ± 85 | 200 | 210 | 156.0 ± 15.3 | 267 | 350 |
| GIPCs | 1428.9 ± 69.4 | - | 1800 | 48 ± 18 | 2000 | - | 236.0 ± 6.7 | 501 | 600 |
| LCBs | 6.4 ± 0.4 | 200 | - | 30 ± 6 | - | 5 | 3.8 ± 0.9 | - | 10 |
| Total | 1818.2 ± 91.4 | 338 | 2115 | 1906 ± 81 | 2590 | 270 | 425.0 ± 20.4 | 782 | 1015 |

[1]These figures were previously published in nmol/g fresh wild type, which is approximately one-tenth the amount in nmol/g dry wt.

[2]CER; non-hydroxy ceramide, HCER; α-hydroxy ceramide, GCER; glucosyl ceramide, GIPC; glucosyl inositol phosphoryl ceramide, LCB; long-chain bases.

The data are expressed as arithmetic mean ± standard error of the mean.

**Supplemental Table S4.** Primers used in this study

| **Primer** | **Sequence** | **Purpose** |
|---|---|---|
| LP_Sail_794_B01 | TCCCAAAAGCAATTCCTCTTAC | T-DNA genotyping |
| RP_Sail_794_B01 | CGTCATAGCTAAGAGGAGGGG | T-DNA genotyping |
| LP_Salk_042166 | TGATATGGAAACAGAGCTGCC | T-DNA genotyping |
| RP_Salk_042166 | TTCCCCACATTTAGCATCATC | T-DNA genotyping |
| LP_CS1011117 | GAATTCAACAACCACAAACCG | T-DNA genotyping |
| RP_CS1011117 | CTGGATCTGAACGAAGACAGC | T-DNA genotyping |
| SPHK1_RT-F | AGAAATGAAAGGCCCATTTG | RT-PCR |
| SPHK1_RT-R | ACAATCAAATCCAGGAAGCC | RT-PCR |
| GCS_RT-F | CATTGGGAAGAAATTTGGCT | RT-PCR |
| GCS_RT-R | ATCACCAACATCGCAAAGAA | RT-PCR |
| PdBG2-RT-F | CACTGATGCAGGGAGAGTCA | RT-PCR |
| PdBG2-RT-R | AGTGGTGGTGACTGTAGCAA | RT-PCR |
| VAMP721-ATTB1-F | AAAAAGCAGGCTTTATGGCGCAACAATCGTTGA | Cloning |
| VAMP721-ATTB2-R | AGAAAGCTGGGTTACACTTAAACCCATGGCAA | Cloning |
| PDLP1_CDS-F | AAAAAGCAGGCTTTATGAAACTCACCTATCAAT | Cloning |
| PDLP1_CDS-NSC-R | AGAAAGCTGGGTTATAAGCATCATATTTATTA | Cloning |
| ATTB1-F | ACAGTTTGTACAAAAAAGCAGGCT | Cloning |
| ATTB2-R | ACCACTTTGTACAAGAAAGCTGGGT | Cloning |
| SP-CB1_ATTB1-F | AAAAAGCAGGCTTTATGGCTGCTCTGGTGCTTT | Cloning |
| SP-CB1-R | GTTCTTCTCCTTTACTCATGGCACCTACAAAGACC | Cloning |
| GFP-CB1-F | TGTGGTCTTTGTAGGTGCCATGAGTAAAGGAGAAG | Cloning |
| GFP-CB1-R | TCTTACACACACACCATGGTGGTGGTGGTGGTGGTG | Cloning |
| CDS+CB1-F | CACCACCACCACCACCACTCATGGTGTGTGTGTAAGA | Cloning |
| CDS+CB1-R | AGAAAGCTGGGTTTTAGAGCATCAGGAAAGAGCAG | Cloning |
| GFP-BG2-R | TCAGAAATTGTTTACCTTGGTGGTGGTGGTGGTGGTG | Cloning |
| CDS-BG2-F | CACCACCACCACCACCACCAAGGTAAACAATTTCTGA | Cloning |
| CDS-BG2-R | AGAAAGCTGGGTTCTACAAGAATACTAAGGCAATG | Cloning |
| ACTIN2-RT-F | AGACCAGCTCTTCCATCGAG | RT-PCR |
| ACTIN2-RT-R | GCAGCTTCCATTCCCACAAA | RT-PCR |
| LBb1.3 | ATTTTGCCGATTTCGGAAC | T-DNA genotyping |
| LB1 | GCCTTTTCAGAAATGGATAAATAGCCTTGCTTCC | T-DNA genotyping |
